# Spike protein fusion loop controls SARS-CoV-2 fusogenicity and infectivity

**DOI:** 10.1101/2020.07.07.191973

**Authors:** Debnath Pal

**Author notes:** Debnath Pal, **Email:**. **Author Contributions** DP did all the work and wrote the paper.

## Abstract

Compared to the other human coronaviruses, SARS-CoV-2 has a higher reproductive number that is driving the COVID-19 pandemic. The high transmission of SARS-CoV-2 has been attributed to environmental, immunological, and molecular factors. The Spike protein is the foremost molecular factor responsible for virus fusion, entry and spread in the host, and thus holds clues for the rapid viral spread. The dense glycosylation of Spike, its high affinity of binding to the human ACE2 receptor, and the efficient priming by cleavage have already been proposed for driving efficient virus-host entry, but these do not explain its unusually high transmission rate. I have investigated the Spike from six β-coronaviruses, including the SARS-CoV-2, and find that their surface-exposed fusion peptides constituting the defined fusion loop are spatially organized contiguous to each other to work synergistically for triggering the virus-host membrane fusion process. The architecture of the Spike quaternary structure ensures the participation of the fusion peptides in the initiation of the host membrane contact for the virus fusion process. The SARS-CoV-2 fusion peptides have unique physicochemical properties, accrued in part from the presence of consecutive prolines that impart backbone rigidity which aids the virus fusogenicity. The specific contribution of these prolines shows significantly diminished fusogenicity *in vitro* and associated pathogenesis *in vivo,* inferred from comparative studies of their deletion-mutant in a fellow murine β-coronavirus MHV-A59. The priming of the Spike by its cleavage and subsequent fusogenic conformational transition steered by the fusion loop may be critical for the SARS-CoV-2 spread.

**Significance Statement:** The three proximal fusion peptides constituting the fusion loop in Spike protein are the membranotropic segments most suitable for engaging the host membrane surface for its disruption. Spike’s unique quaternary structure architecture drives the fusion peptides to initiate the protein host membrane contact. The SARS-CoV-2 Spike trimer surface is relatively more hydrophobic among other human coronavirus Spikes, including the fusion peptides that are structurally more rigid owing to the presence of consecutive prolines, aromatic/hydrophobic clusters, a stretch of consecutive β-branched amino acids, and the hydrogen bonds. The synergy accrued from the location of the fusion peptides, their physicochemical features, and the fusogenic conformational transition appears to drive the virus fusion process and may explain the high spread of the SARS-CoV-2.

## Introduction

The novel Coronavirus Disease 2019 (COVID-19) caused by SARS-CoV-2, Severe Acute Respiratory Syndrome (SARS) by SARS-CoV, and Middle Eastern Respiratory Syndrome (MERS) by MERS-CoV, induce severe acute respiratory distress in patients. Though these diseases share similar clinical and pathological features, COVID-19 differs in overlapping yet distinct phases of infection (1, 2). The degree of infectivity is significantly high in SARS-CoV-2, and far more aggressive, as evidenced by the current global pandemic. This can be quantified by the preliminary reproductive number (R_0_) of COVID-19 (2.0-2.5), which is higher than the R_0_ of SARS (1.7-1.9) and far higher than that of MERS (<1) (3). The significant difference in R_0_ may accrue due to environmental, immunological or molecular reasons. COVID-19 transmission has been attributed to the long life of SARS-CoV-2 outside the host as it increases the chances of infection through cross-contamination by contact in the population (4). The large distance distribution of the SARS-CoV-2 particles from the infected person due to activities like sneezing and coughing (5), and the tiny size of the virus droplets may be more efficient in penetrating deeply into the pulmonary system to allow rapid spread of the disease (6). However, the SARS-CoV has high genomic similarity with the SARS-CoV-2, and one would have expected it to have similar transmission behavior and R_0_, which is evidently not the case. For that matter, the environmental spread of other viruses should have been far more widespread than coronaviruses, given that the coronaviruses have the largest RNA viral genomes and therefore the largest particle size and consequently higher aerosol size compared to many other viruses. This is again not that we observe in practice. Another possibility of high viral spread may accrue from intense viral shedding, where SAR-CoV-2 has succeeded early on in rapid viral replication and cell-to-cell spread before the onset of acute inflammatory response. Here the extent of the viral replication is dependent on the immune containment-response, but given that MERS has shown 34% case fatality compared to 9.5% for SARS-CoV and only 2.3% for SARS-CoV-2 thus far (3), one can argue that the R_0_ values should also have been in the same order, which is evidently not the case. This suggests that neither environmental nor immunological response suitably explains the higher R_0_ of SARS-CoV-2, suggesting that the key reason may be molecular. The Spike glycoprotein that protrudes |150 Å out of the 500-2000 Å diameter coronavirus envelope is the most suitable molecule for making the first contact with the host cell, and is, therefore, a key molecular factor that determines virus fusion, entry and spread in the host, and thus holds clues for the rapid spread of SARS-CoV-2.

Spike (S) is a key well-studied gene among all coronaviruses that determines virus infectivity due to its established role in initiating virus-host attachment, fusion, entry, and cell-to-cell fusion post replication (7, 8). The expressed protein executes its function through a trimeric quaternary structure, with each monomer segmented into the S1 and S2 domains, where S1 consists of the N-terminal domain (NTD) and the C-terminal domain (CTD), while the S2 (also known as the fusion domain) consists of the fusion peptides (FPs), central helix (CH), heptad repeat regions HR1 and HR2, and a transmembrane domain (TM) followed by the cytoplasmic tail (CP) (Fig. 1A). The NTD or CTD embeds the receptor-binding domain (RBD) depending on the individual virus. In SARS-CoV-2 the CTD embeds the RBD. The RBD recognizes the host-cell and mediates the virus-host attachment through interaction with a receptor, while the S2 domain drives the viral entry into the host cell and cell-to-cell fusion by catalyzing the virus-host membrane fusion process. The TM anchors the protein in the viral envelope, while the FPs contribute to the trigger that drives the virus-host membrane fusion process. A transition pathway has been proposed to proceed through a pre-hairpin intermediate structure followed by pre-bundle hemifusion membrane-associated structure, bundle structure, and the eventual post-fusion structure (9). It is remarkable that despite their similarity in function, β-coronavirus Spike proteins have widely diverged, with the RBD diverging further (21%-61% pairwise sequence identity), explained in part by the diversity of its host receptor targets like the human ACE2 for SARS-CoV and SARS-CoV-2, DPP4 for MERS-CoV, and 9-O-Ac-Sia for HCoV-OC43 and HCoV-HKU1. The S2 domain has also diverged, but comparatively less with >33% overall pairwise sequence identity – the heptad repeat regions being the most conserved (Fig. 1B). It is pertinent to ask what features of Spike sequence and structure determine the virus fusogenicity and infectivity.

**Figure 1.**
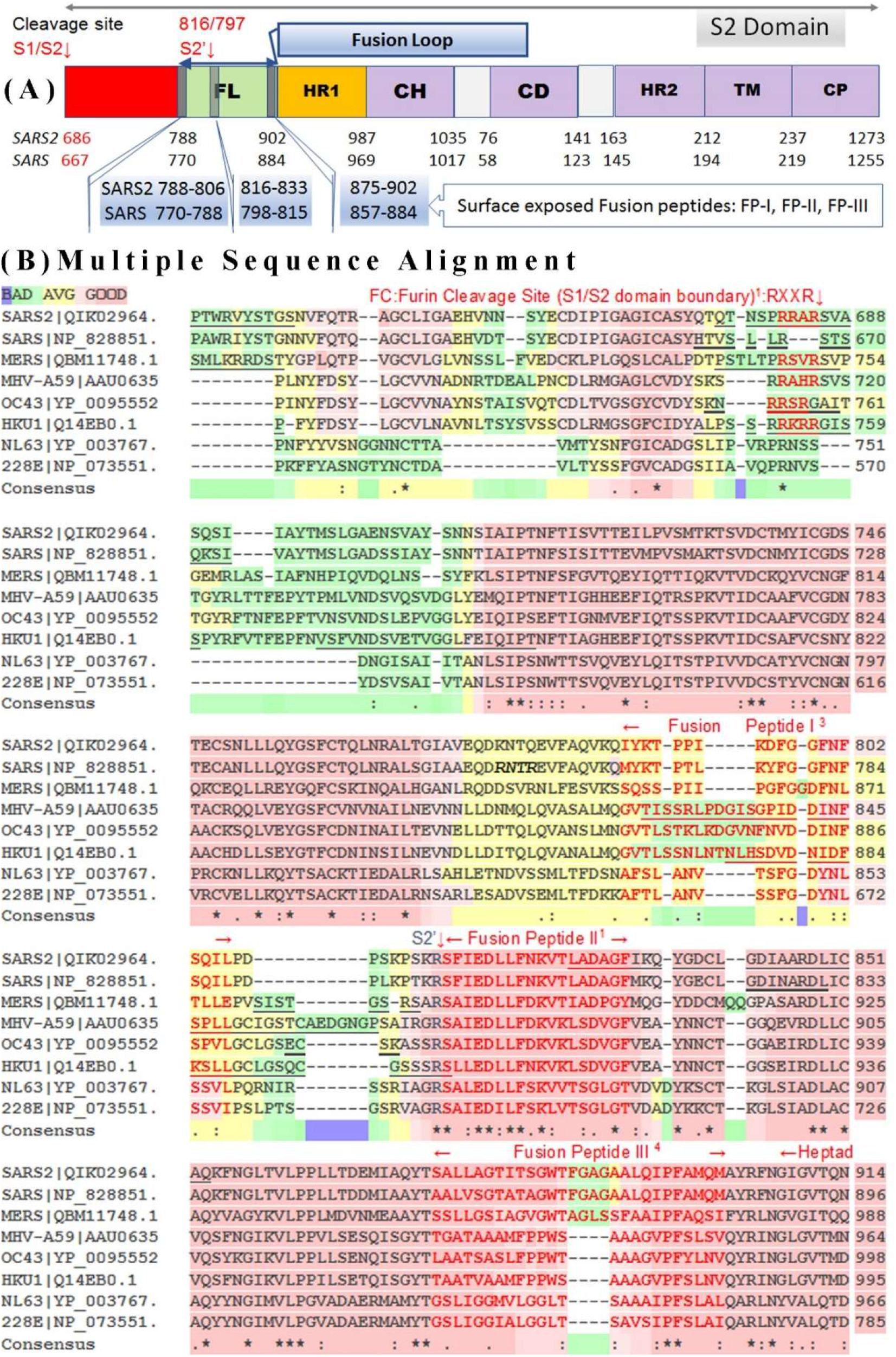
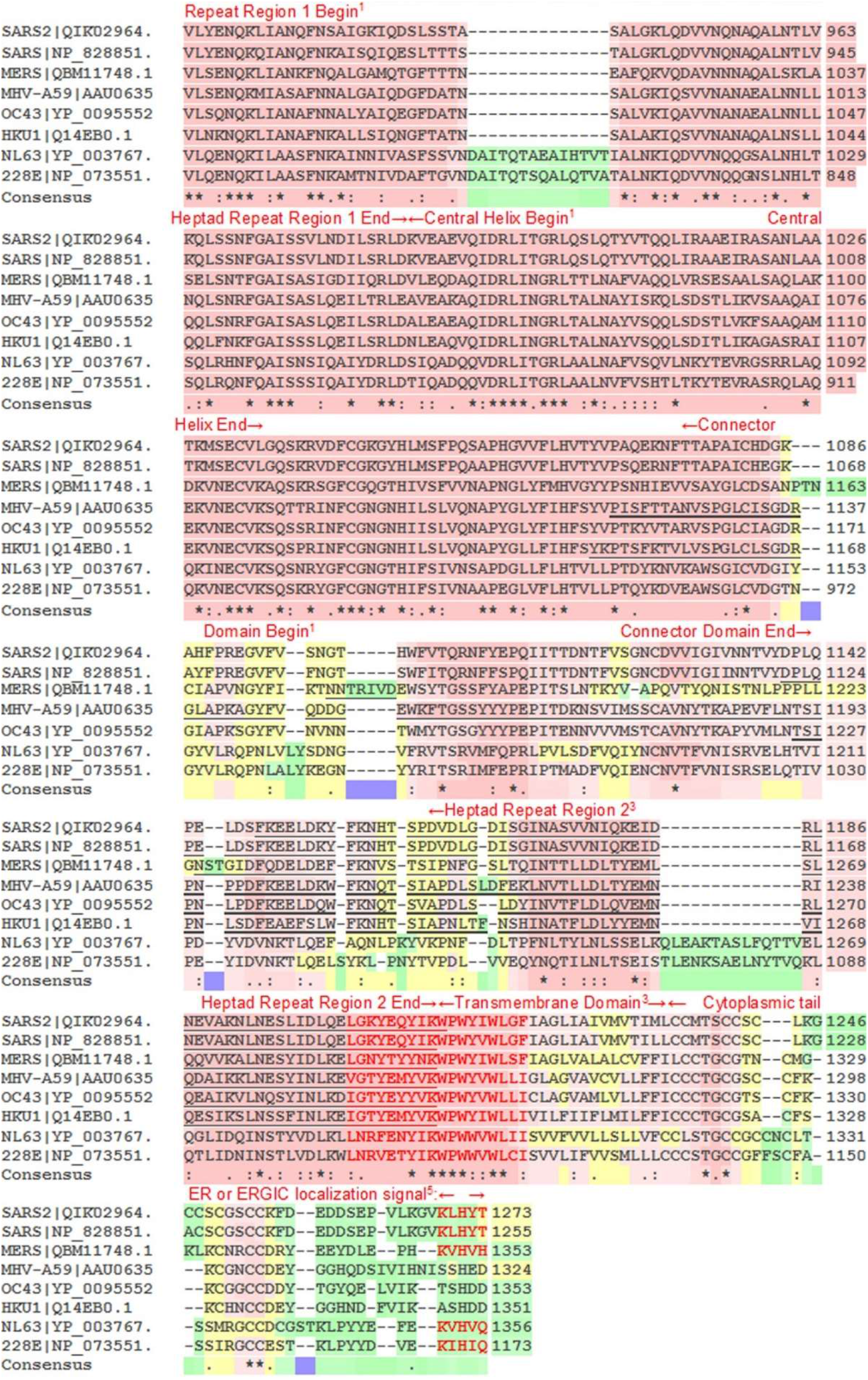
The S2 domain organization of SARS-CoV-2 Spike glycoprotein and its sequence comparison against seven other coronaviruses. (A) The denomination of the various S2 domain segments and their positions. The labels are as follows, FL: Fusion Loop, HR1: Hepatad Repeat 1, CH: Central Helix, CD: Connector Domain, HR2: Heptad Repeat 2, TM: Transmembrane Domain, CP: Cytoplasmic tail. The span of the segments can be estimated from the position markers indicated for SAR-CoV-2 (SARS2) and SARS-CoV (SARS). (B) Multiple sequence alignment (MSA) of the S2 domain of Spike protein from eight coronaviruses. The segments of the S2 domain are also marked by position markers. Residues <u>underlined</u> in the MSA have no coordinates in the structure file and are deemed flexible regions. The PDB files used to mark the same are SARS-CoV-2: 6VXX, SARS-CoV:5XLR, MERS-CoV: 6Q04, MHV-A59: 3JCL, HCoV-OC43: 6NZK, HCoV-HKU1: 5I08. The sequences for HCoV-NL63 and HCoV-2289E are not marked because they belong to α-coronaviruses; whereas, all others are from β-coronaviruses. The RED marked region spanning Heptad repeat region 2 C-terminal and trans-membrane domain N-terminal region is also called the aromatic domain or the FP-IV. The MSA of the full-length protein and further annotation details can be found in Fig. S1.

Densely glycosylated Spike protein has been suggested as the prime reason for the high SARS-CoV-2 infectivity (7). This extensive glycosylation of the S2 domain is driven by an intracellular N-terminal signaling peptide for transport and retention in the endoplasmic reticulum (see Fig. 1B Multiple Sequence Alignment bottom panel). However, this signal sequence is absent in a fellow murine β-coronavirus MHV-A59 Spike protein (10), indicating limited glycosylation. Yet, MHV-A59 aggressively infects the mouse liver and brain. Upon intracranial inoculation in mice, it can cause acute stage meningoencephalitis and myelitis, chronic stage demyelination, and axonal loss (11-13). It infects the neurons profusely and can spread from neuron to neuron. Its propagation from grey matter neuron to white matter and release at the nerve ends to infect the oligodendrocytes by cell-to-cell fusion (12) are robust mechanisms to evade immune responses and induce chronic stage progressive neuroinflammatory demyelination concurrent with axonal loss in the absence of functional virions (12, 14). Therefore, high infectivity of the MHV-A59 Spike does not appear to be contingent on glycosylation and one can argue the same for SARS-CoV-2 Spike, where its glycosylation may only marginally raise the basal fusion efficiency. Surface glycosylation thus may not be a contributing factor to host cell binding, although the successive virus-to-cell and cell-to-cell fusion may all-together play an important role in higher virus infectivity.

It has been suggested that enhanced virus-to-cell infection can be propelled by the increased number of hydrogen-bonded contacts between SARS-CoV-2 RBD and ACE2 receptor leading to higher affinity and improved host targeting compared to the SARS-CoV (7, 15). However, a significantly higher affinity between ACE2 and SARS-CoV-2 has not been experimentally corroborated (16). Besides, such a proposition is weak because the RBDs in all HCoVs are diverse, including SARS-CoV, where the minimal RBD (318 to 510 residues) (17) shares only 74% sequence identity with SARS-CoV-2 (Fig. S1). Also, the SARS-CoV-2 Spike may interact with other receptors such as DC-SIGN and DC-SIGNR as in SARS-CoV to increase tropism (18) and viral spread. Therefore, there is no direct consequence of ACE2 recognition with infectivity unless a virus entry can be realized; however, when RBDs interact with ACE2 in large numbers during the acute stage of the infection, they may modulate the host immune response by downregulating hydrolysis of the pro-inflammatory angiotensin II to anti-inflammatory angiotensin 1-7 in the reninangiotensin signaling pathway (19). This can alter the immune response and increase infectivity. But such effects can manifest only beyond the early stage of the infection, and for that to happen the efficiency of viral entry is the rate-limiting step.

The cleavage of the Spike protein is said to prime it for the efficient virus-host membrane fusion process. How essential is this for virus fusogenicity and infectivity is an important consideration. The Spike cleavage potentially removes any *in situ* covalent and noncovalent constraints that the S1 domain may impose on the S2 domain impeding its conformational transition that facilitates the virus entry. It has been proposed that SARS-CoV-2 Spike is preactivated by cleavage at the S1/S2 site when it is packaged inside the host, and the S2’ site is cleaved when the Spike gets attached to the host receptor, which makes the priming process very efficient (20). However, a comparison of the S1/S2 cleavage signal sequence …RXXR… shows that SARS-CoV-2 “…RRARS…” Furin recognition site is similar to MHV-A59 Spike’s “…RRAHR…”, and others like MERS, HCoV-OC43 and HCoV-HKU1 Spike have a conserved motif sequence as well (Fig. 1B). The cleavage site signal at S2’ embedding a single Arginine is highly conserved across all HCoVs. Therefore, the efficient priming advantage available to SAR-CoV-2 Spike is equally present for MHV-A59, MERS, HCoV-OC43, and HCoV-HKU1 Spike. In contrast, the canonical S1/S2 cleavage recognition sequence is missing in SARS-CoV with only a single Arginine present there. A regular cleavage at this site has not been reported, and cleavage by trypsin has been shown to activate the virus independent of the pH due to the presence of a single Arginine. The importance of this region has been aptly corroborated by S2’ site cleavage studies in SARS-CoV (21). Besides, it is also possible for Spike to be activated by the low pH environment through protonation of residues if it internalizes in the endosome post interaction with the host receptor. In contrast, fusion processes are known to happen in MHV-A59 Spike without cleavage as well (22). Therefore, based on the similarity SARS-CoV-2 Spike with others, one can argue that it is competent to access multiple pathways for priming that enhances its infection capability, including the possibility that it can infect without a cleavage as well – though these advantages are not unique.

Among all the components of the fusion apparatus in the Spike S2 domain, the FPs are the least studied although they have been suggested to contribute to the trigger that drives the virus-host fusion process by initiating the protein-host membrane contacts. Limited experimental information available shows mutation in FP of SARS-CoV Spike can significantly perturb the fusion efficiency (23) as much as >70% (24). Most studies of the FP regions have used synthetic peptides in a fusion assay system to understand their membrane perturbing capabilities (25-28), and how Ca^+2^ ions may interact with these peptides to modulate fusion (9). Interestingly, the FPs also contains a central proline (29) in several viruses such as the Avian Sarcoma/Leucosis virus (30), Ebola virus (31), Vesicular Stomatitis virus (32), and Hepatitis C virus (33), where its important role has been investigated through mutation studies. The location of coronavirus FPs proximal to the N-terminal of the S2 domain is reminiscent of FPs from HIV-1, influenza virus, and paramyxoviruses. They have been suggested to be located at the head a pre-hairpin intermediate structure (34) predicted for the current model of class I viral fusion proteins.

Although the FPs are believed to be the early initiators of protein and host membrane contact, there is still no consensus on their location. For example, the fusion peptide for SARS-CoV-2 Spike has been cited at 788-806 position by Xia et al. (8) compared to 816-833 by Wrap et al. (7). When inferred from alignment to SARS-CoV Spike, two additional FP segments at 875-902 position and 1203-1220 can be proposed based on experimental studies by Ou et al. (35) and Guillén et al. (26), respectively. In reality, all four Spike fusion domain segments (FP-I to FP-IV; Fig. 1B) mentioned can be identified by a simple window-based analysis using interfacial hydrophobicity scales such as from Wimley and White (36). Given that FP-1 to FP-III are contiguous to each other in the sequence, the whole segment spanning the beginning of FP-1 to the end of FP-III can be termed together as the “fusion loop” (Fig. 1A). But how these FPs in the loop can act synergistically to rapidly trigger the membrane fusion process is what we explore further in this study.

## Results

### Spike receptor binding and the fusion loop

To understand the synergy of the FPs in the trigger process, it is important to understand the structure of the Spike protein in the proper context. The FP-I to FP-III are surface exposed and are contiguous to each other in space as seen from the three-dimensional structures of the Spike fusion domain (Fig. 2, cartoon diagrams). While FP-I to FP-III are always surface exposed in the full-length, the FP-IV is deeply buried and interface the virus membrane. (The location of FP-IV although not available from the three-dimensional structure can be inferred from the primary structure and the location of the transmembrane domain (Fig. 1B)). Therefore, while FP-I to FP-III are always available for early contacts with the host membrane, FP-IV can participate in the process post the conformational transition which may expose it for interaction with the host membrane. To understand what guarantees the FP surfaces to make the initial protein-host membrane contact, one must look at the possible modes of virus-host attachment mediated by the Spike. For this, we propose a new contact initiation model, where there is no requirement of a fusion peptide to be at the N-terminal of the conformationally transformed pre-hairpin intermediate (9). To understand the model, let us consider the different options for host receptor binding. For example, if all three RBDs in the trimeric structure find the host receptors, it can attain a tripod binding mode (Fig. 3A). A recent structure of the trimeric Spike complexed with a host receptor reveals the precise geometry of CEACAM1 binding the RBD of Spike from MHV-A59 (37). However, a tripod binding requires receptor molecules on the host membrane to be pre-available in a specific arrangement. High expression of receptor molecules on the host cell surface is expected to increase the probability of tripod binding, but there is no existing information whether such a precise arrangement is present on the host surface suitable for interaction with the trimeric RBD. Moreover, the membrane bilayer structure does not contain any feature that can direct such regular host receptor arrangement. It may be noted that only in a tripod arrangement, the N-terminal segment of the S2 domain will interact early during the protein-membrane contact post intermediate structure formation, and in such a case the pre-hairpin/pre-bundle helices of the fusion domain are expected to interact head-on with the host membrane (9). However, FP-I and FP-II are located at the middle of the cylinder-like Spike S2 structure (Fig. 3A) and an intermediate formation through conformational transition is needed to place them near the head of the cylinder. Here, FP1 is expected to make the early host-membrane contact if the cleavage is at the S1/S2 site and FP-II if the cleavage is at the S2’ site (see Fig. 1A). The FP-III region has limited scope to make any early contact due to its farthest position from the host membrane surface. In tripod binding mode the virus membrane is still ~150Å away, and the three HR2 regions need to fold back and bind to the hydrophobic grooves of the HR1 trimer in an antiparallel manner to bridge this gap and form a hemifusion structure with the host membrane (9).

**Figure 2.**
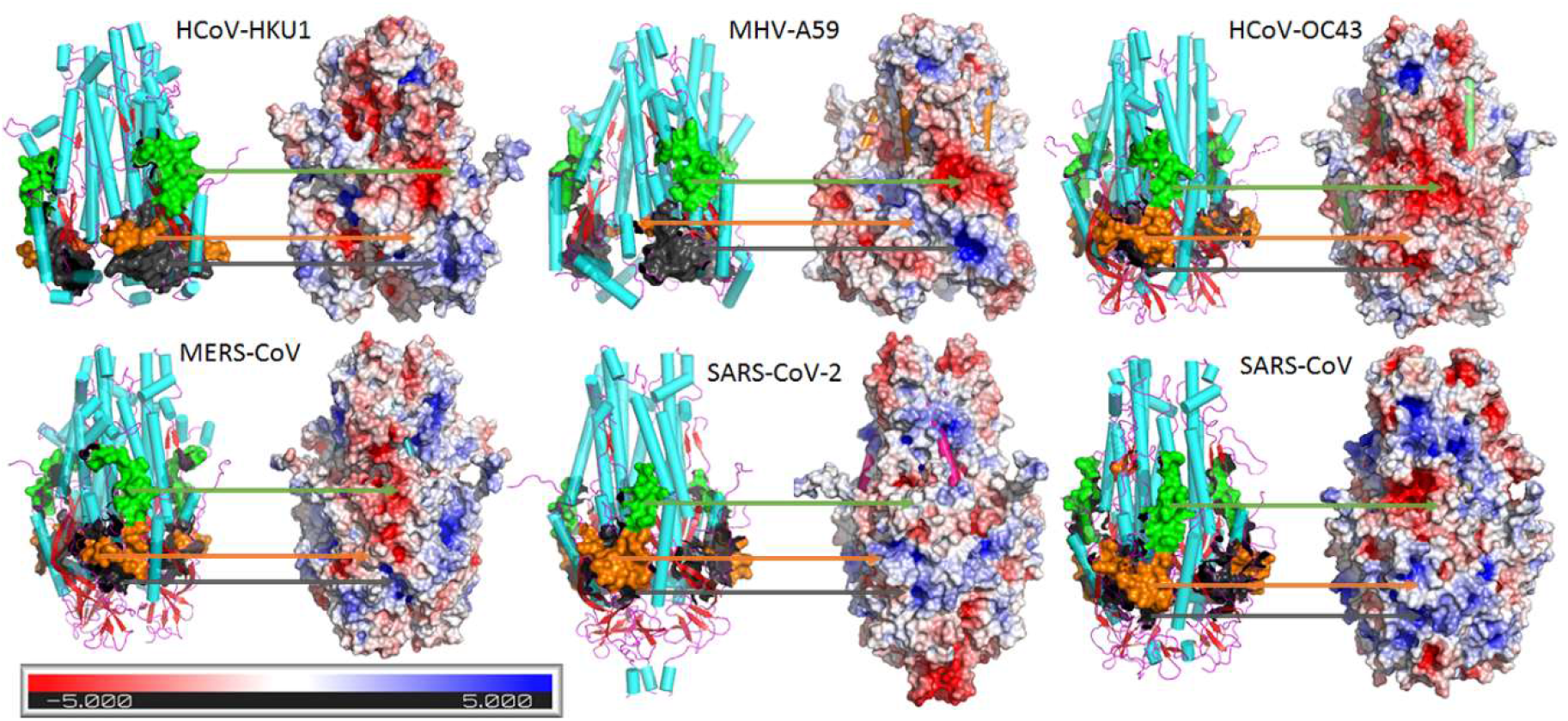
Cartoon diagram showing the fusion peptide regions FP-I (orange), FP-II (green), and FP-III (dark grey) on the trimeric fusion domain structure of Spike from six viruses. The electrostatic surface of the fusion domain is shown adjacent to each structure and the corresponding fusion peptide regions are marked by an arrow. The orientation of the cartoon structure and the electrostatic surface are aligned among themselves. Note that one face of the electrostatic surface that is visible is repeated on the other side due to the symmetry arising out of the trimeric quaternary structure. The fusion domain structures are shown in cyan for helices, strands in red, and loops in magenta. Please refer to Fig. 1 for the sequence alignment and its legend for the PDB ID of files used to draw the structures. Note that a part of the FP-I surface is absent for MHV-A59 and HCoV-HKU1 due to unavailable atom coordinates in the PDB file. The same is true for a small section of FP-II from SARS-CoV-2. For residue details, please refer to Fig. 4. All FP-I regions have at least one Asn-linked site, only SARS-CoV-2, SARS-CoV, and HCoV-HKU1 has a site in FP-II, and HCoV-OC43 and HCoV-HKU1 in FP-III. Contiguous to the FP-III are the heptad repeat regions where the glycosylation and sequence conservation among Spike is the highest. Spike from MHV-A59 shares a 64-66% overall sequence identity with HCoV-HKU1 and OC43, 71-76% identity with the corresponding S2 domains. Similar match with SARS-CoV, MERS-CoV, and SARS-CoV-2 are at <30% pairwise sequence identity.

**Fig. 3.**
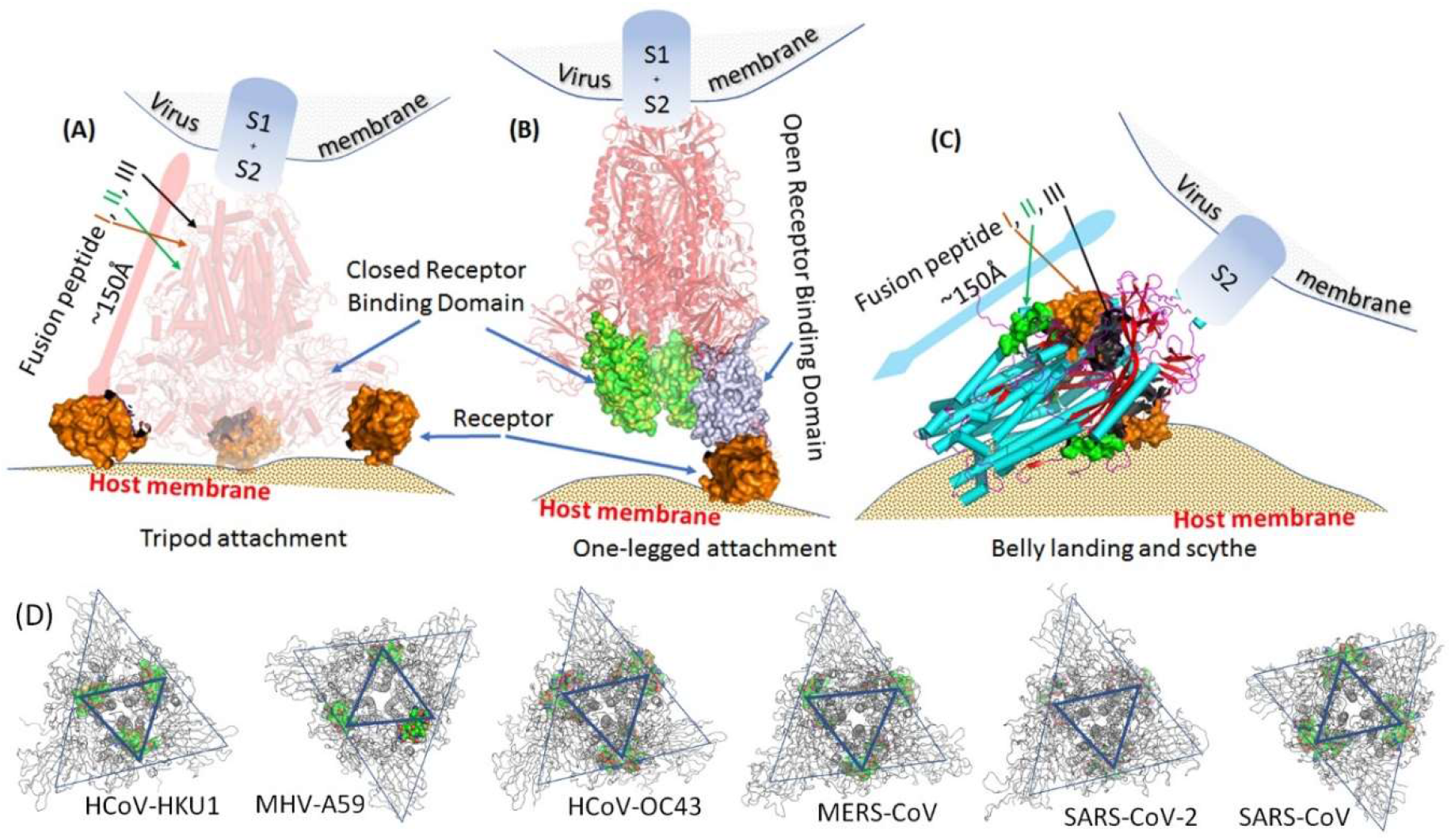
Schematic diagrams explaining the mode of binding of Spike protein to the host receptor and its putative orientation relative to the host membrane surface during the virushost attachment. (A). A cartoon diagram created using PDB ID: 6VSJ where the Spike protein from MHV-A59 is in complex with CEACAM1 receptor from mouse. Since all three subunits from the Spike RBD attach to the receptor, it allows the virus to anchor in a tripod mode. The approximate length of the Spike in the longer dimension is indicated along with the FP locations. (B). A cartoon diagram created using PDB ID: 6VSB, where one of the RBDs are shown in an open conformation. A receptor molecule ACE2 has been drawn to show the putative attachment in one-legged mode. Two-legged mode may also be possible similarly. (C). The S2 domain of SARS-CoV-2 Spike protein is shown in a belly-landing orientation. The approximate length of the Spike in the longer dimension is indicated along with the marked FP locations. The S1 domain is likely to be loosely bound and not shown for clarity. (D) A ribbon diagram of the trimeric Spike proteins from the six coronaviruses used in this study. Two triangles are marked on each structure, where the relative locations of the protruding NTDs appear near the outer triangle vertices, and the FP-I, FP-II, and FP-III colored surfaces are located near the inner triangle vertices. The triangles are marked to bring out the relative positions of the NTDs and the FP surfaces. The Spike structure is oriented such that the RBD appears closest to the eye, followed by the NTD and then the FPs.

### The Contact Initiation Model - Spike fusion peptide trigger

If only one or two RBDs bind the receptor, the vertical anchoring of the Spike fusion domain relative to the host surface lacks the third anchor rendering the vertical orientation unstable and unfeasible. Also, a recent trimeric structure of SARS-CoV-2 Spike has shown a single RBD to be in the open conformation (7), where it is swiveled away from its core structure originally interacting with the NTD and the fusion domain. In such a state, the interaction of the open RBD with the host (Fig. 3B) is not expected to stabilize the Spike anchoring in any specific orientation relative to the host due to the weak interaction with the fusion domain. Here a post-cleaved S2 domain is expected to interact side-on to the membrane surface through a “belly” landing to trigger the fusion process (Fig. 3C). Such a process would be sterically facile if there are no other Spike in the vicinity on the virus surface. It is to be noted that the shape of the trimeric S2 domain is not a proper cylinder, but with a bulge in the mid-segment which we call a “belly”. The FP-II and FP-I surfaces are located at the crest of this bulge, such that it is able the make the initial contact with the host membrane. The structural constraints that guarantee the “belly” landing can be understood from the overall geometry of the Spike (Fig. 3D). It is to be noted that the most stable and eventual landing posture of an object having an uneven surface will be the one that guarantees the largest surface area of contact with the host landing surface. The Spike architecture is such where the relative location of the NTDs approximately form three vertices of a triangle, while the FP-I to F-III are located midpoint of the sides, forming the vertices of an inner triangle. Based on the maximum landing surface premise, if we consider the receptor attachment to be in two RBD locations, the S2 landing will be close to the midpoint of the two vertices of the outer triangle coincident to the FP surfaces. Even for one-legged attachment, the contact must always be directed towards the midpoint of the two NTDs because that allows maximum contact surface to be formed where the Spike can stably rest on the host surface. During the contact, the membranotropic segment of the fusion loop spanning the FPs is expected to engage the host membrane during the fusogenic conformational change. In this case, the FP sites are proximal to the virus membrane surface such that hemifusion membrane structures can be initiated early in comparison to the tripod-binding mode which requires an intermediate pre-hairpin structure to be formed. It is also to be noted that weak RBD binding to S2 domain or disintegration of the Spike trimer post tripod binding can mimic the one- or two-legged binding mode. For additional lucidity, the structural basis of the Contact Initiation model is further explained pictorially in a lay manner in Fig. S2.

### SARS-CoV-2 fusion peptide distinctions

The physicochemical property of the fusion loop and the synergy of the FPs therein is critical to the rapid initiation and transition to the hemifusion stage of the membrane fusion process. The proposed Contact Initiation Model ensures that the S2 domain lies on the belly contacting the host surface during the conformational transition. Since the exerted force during the conformation transition is tangential to the host surface, the orthogonal frictional forces may allow an efficient scything action based on the physiochemical nature of the contact engaged by the fusion loop. The nature of the initial surfaces of the fusion loop can be obtained from the electrostatic potential of the FP-I to FP-III surface patches (38) (Fig. 2, surface diagrams). Among the six Spike proteins considered in this study, SARS-CoV-2 has the most hydrophobic/neutral electrostatic FP patches (WHITE colored) most suitable for membrane disruption. It also has the least amount of highly negative electrostatic surface (RED colored patches) which repel the attachment of the S2 domain to the host membrane due to the repelling hydrophilic surface of the outer membrane composed of negatively charged fatty acid groups. On the other hand, a positive electrostatic surface (BLUE colored patches) may allow protein surface to tightly attach to the membrane exterior through charge attraction. The presence of an interfacial ion like the Ca^+2^ can cap the negative charge at the FP surface to assist fusion trigger - consistent with the membrane charge compatibility requirements. The disruption efficiency of surface patches is, therefore, likely to be highest for the hydrophobic/neutral, followed by the positive and negative electrostatic potential, unless modulated by an interfacial cation like the Ca^2+^. One may argue that large patches of negative electrostatic surface potential (RED color) in the HCoV-HKU1 and HCoV-OC43 Spike fusion domain may explain the mild nature of those viruses.

Aside from the electrostatics, the physical rigidity of the fusion loop in Spike is of prime importance for the virus fusogenicity. This is because the Spike fusion domain being metastable, any local alteration of rigidity has global implications for the molecule. This has been alluded to by the mutation studies on the fusion peptide central prolines (30-33), but its criticality was recently revealed from our comprehensive studies on centrally located consecutive prolines in MHV-A59 Spike fusion peptide (39). Proline being an imino-acid with unique structure, has restricted torsional freedom, which in turn restricts the torsional freedom of the protein backbone where it is located (40). When two consecutive prolines are located, the rigidity of the protein backbone is further enhanced. Among the six Spike proteins in our study, three have a single central proline in FP-I, and two centrally located consecutive prolines occur in SARS-CoV-2 Spike FP-1. There are no central prolines in FP-II and a single conserved central proline in FP-IV, while consecutive prolines exist at a central location in HCoV-HKU1, HCoV-OC43, and MHV-A59 FP-III Spikes. To understand the intrinsic flexibility/rigidity of the surface exposed FPs, we set up 500 ns molecular dynamics simulations (Fig. 4). The starting structures of all the FP-I fragments are devoid of stable secondary structures such as helices or sheets and is dominated by turns, and irregular structures indicating that the region prefers to be in a loop conformation. This is true even for the MHV-A59 Spike, which is flexible as experimentally evidenced by the lack of coordinate from the electron density map for complete FP-I in the PDB file: 3JCL. In comparison, FP-IIs in all cases are dominated by helical conformation, with the N-terminal FP segments always in a helical state. This is true for FP-III region fragments as well. Given an irregular structure, the effect of proline is more dramatic on the FP-I region in contrast to FPs from FP-II and FP-III which are already stabilized by hydrogen bonds in helices. For all FPs in the FP-I region, although local secondary structures are induced during simulation, the lowest root-mean-square fluctuation (RMSF) for a residue is achieved only by the SARS-CoV-2 consecutive proline containing FP. This can also be confirmed from the energy landscape plot created from the molecular dynamics simulation trajectory (Fig. S3) where it has the most compact single conformational well among all the FP-Is. If we look at the FP-III regions where other consecutive proline containing FPs exist, HCoV-OC43 has the lowest residue RMSF at the double prolines; SARS-CoV-2 FP-III segment does not have any central proline, but it contains a unique “Thr-Ile-Thr” segment constituted of three consecutive β-branched side-chain residues that can impart substantial rigidity based on steric considerations (40) if not as much as the consecutive prolines. This can be again confirmed from the energy landscape plot which shows a single compact conformational well in contrast to the other FP-IIIs (Fig. S3). The observations are consistent with our previous comparative molecular dynamics studies (39) on MHV-A59 FP-III and single central proline containing FP-III from another parental non-neurotropic strain of murine hepatitis virus (MHV), MHV-2, where we found the FP-III from the former became more rigid than the latter in methanolic conditions compared to water. Additionally, NMR studies on the MHV-A59 FP-III fragment revealed its unique ability to form cis-peptide at the central P-P peptide bond (39). A cis-trans isomerization during the membrane fusion process has the potential to expose the hydrophobic residues efficiently around the isomerized-peptide neighborhood, enhancing the fusion trigger potential. While the structural role of proline cannot be discounted, other stabilizing interactions also dominate the FPs, among which the formation of the aromatic/hydrophobic clusters is of relevance to the fusion process (see Fig. S3 for examples). As observed from the packing of the loops, a combination of aromatic and Val/Ile/Leu side chains pack tightly to exclude water. But when these regions become exposed during the conformational transition, they would enhance the hydrophobic interaction of the protein surface with the membrane. In all S2 domains, however, FP-III regions are in part masked by the FP-I segment, as evident from the relative solvent accessible surface area (SASA) values indicated in Fig. 4. The FP-III region can get fully exposed post cleavage at the S2’ site which dislodges the sheathing FP-I segment, or when the site gets progressively exposed during the conformational transition amidst the fusion process.

**Figure 4.**
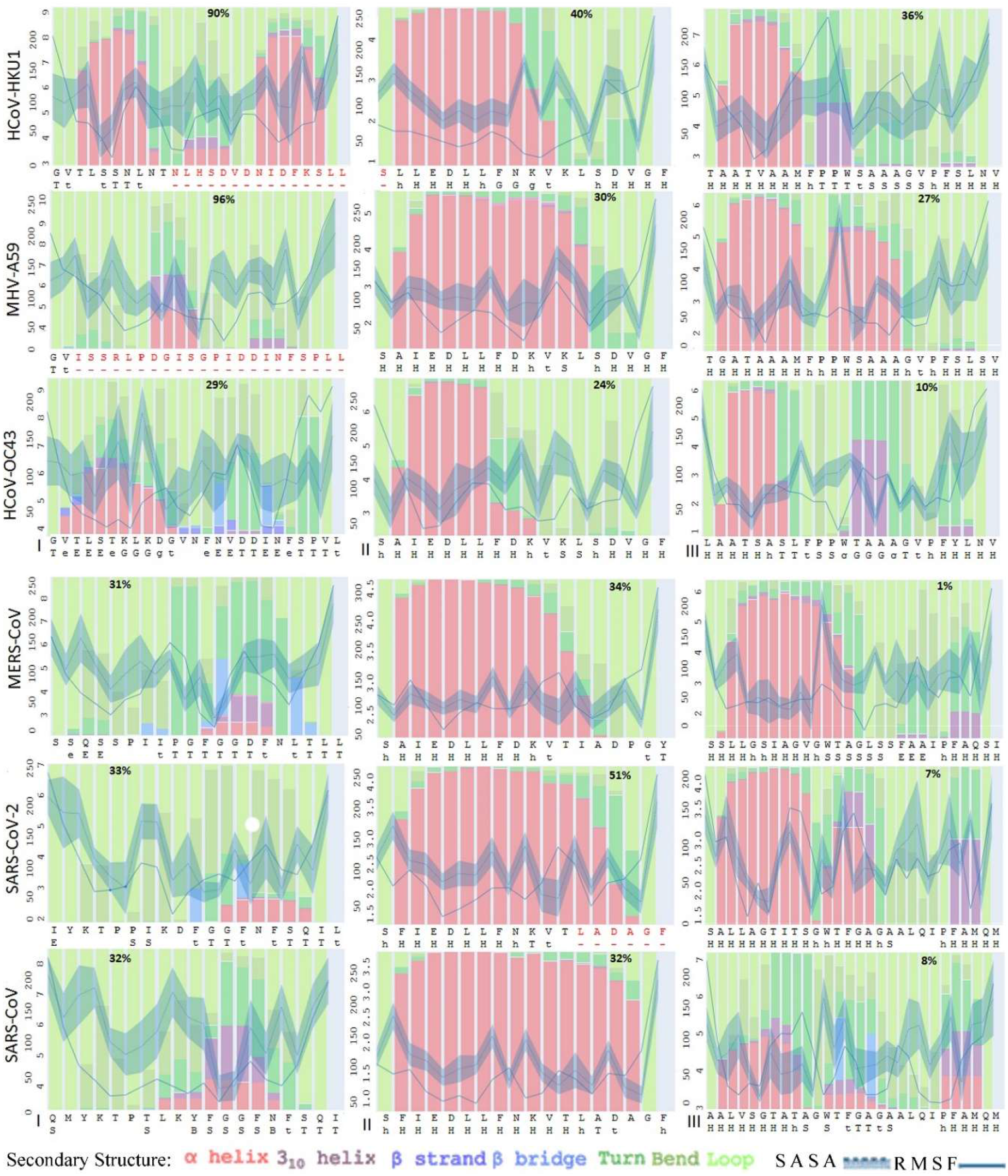
Graphs showing various parameters calculated from the simulation trajectory of the isolated fragments from FP-I, FP-II, and FP-III of six coronaviruses. The bar diagrams are drawn showing the secondary structure prevalence in 500 ns simulation, overlaid by the curves for Solvent Accessible Solvent Area (SASA; Å^2^), and Root-Mean-Square-Fluctuation (RMSF; Å). The Y-axis scales provide values for SASA and RMSF side-by-side. The standard deviation of the SASA in the simulation is indicated by a shaded background around the SASA curve. The X-axis indicates the residues for each FP fragment. The central proline residues are marked in BOLD and those residues for which the atom coordinates are absent in the PDB file are marked in RED. Below the residues, the corresponding secondary structures as indicated as present in the full-length protein. The symbols mean as follows, H:α-helix, h: α-helix termini, G: 3_10_-helix, g: 3_10_-helix termini, B: β-bridge, E: β-strand, e: β-strand termini, T: hydrogen-bonded turn, t: hydrogen-bonded turn termini, S: Bend, and <space>: irregular secondary structure. The first, second, and third columns are indicated by Labels I, II, and III corresponding to FP-I, FP-II, and FP-III in each Spike protein from a given virus, respectively. The relative SASA value of the whole FP fragment in the S2 domain expressed as a percentage is indicated in BOLD on the top part of each plot.

### Experimental corroboration

Additional evidence of “rigidity” imparted by proline being critical for virus fusogenicity, infectivity and pathogenicity can be obtained from our previous in vitro and in vivo studies (39). We generated the Spike containing two consecutive prolines at 938-939 (S-MHV-A59(PP)) in the FP-III and its proline deletion (Δ938) mutant S-MHV-A59(P) from isogenic recombinant strain of MHV-A59, RSA59 (39) Spike protein. The proline deleted mutant only Spike gene construct S-MHV-A59(P) show slower trafficking to the cell surface, and significantly less fusogenicity. Established on differential properties of proline deletion in Spike construct, we generated proline deleted targeted recombinant mutant strain RSA59(P-) which contains S-MHV-A59(P) and compared it with two consecutive proline-containing parental isogenic strain RSA59 containing S-MHV-A59(PP). Proline deleted mutant RSA59(P-) infection in neuronal cell line demonstrated less aggressive and fewer syncytia formation, and one order lower viral titers postinfection *in vitro*. Based on these *in vitro* differential properties of proline deletion in Spike construct we generated proline deleted targeted recombinant mutant strain of RSA59(P-) which contains S-MHV-A59(P) and compared it with two consecutive proline containing parental isogenic strain RSA59 containing S-MHV-A59(PP). The *in vivo* studies in mice parallels the *in vitro* studies demonstrating significantly reduced viral replication and consecutive disease pathologies, like less severity in meningitis, encephalitis, and demyelination, and inability to infect the retina nor induce loss of retinal ganglion cells (41). The non-neurotropic strain MHV-2 sharing 91% genome identity with MHV-A59, and 83% pairwise Spike sequence identity, with a single central proline in FP-III, causes only meningitis and is unable to invade the brain parenchyma. Computational studies of S2 fusion domains of S-MHV-A59(PP) and MHV-2 Spike involving molecular dynamics confirmed the former to be more rigid and containing more residues in the regular secondary structure (39).

### Target for therapy

Our study brings out the importance of the fusion loop region which could be a legitimate target for the design of vaccine or synthetic agents for therapy against COVID-19. The S2 domain serves as a better therapeutic target than S1 due to higher evolutionary conservation, but the few attempts made have mainly focussed attention on the heptad repeat regions (9). Two important features that also make the fusion loop an attractive therapeutic target are its accessibility due to its surface exposure in the full-length Spike, and the relatively high conservation of residues in the fusion loop, especially around the FP-II region which increases its scope as a pancoronavirus target that can cater to future pandemic threats as well. Mimetic peptides can be designed to bind to the fusion loop to inhibit the fusogenic conformational transition of the S2 domain. Impairment of the fusion trigger would have a direct bearing on the fusogenicity of the virus and contribute to the reduction of lung invasion and damage that clinically results in acute pneumonia. Systematic studies can identify the minimal motif in the fusion loop serving as the fusogenic determinant to improve our selection of a potential therapeutic target to prevent cell-to-cell fusion and subsequent pathogenesis.

## Discussion

The interplay of the outlined physicochemical features determines the virus-entry process to become more efficient. The local and global stability of the S2 domain is important. Since the S2 domain undergoes a conformational transition, local stability means reasonably rigid moving parts, and global stability means a well-defined conformational transition pathway from the metastable to the stable state. This local and global stability requirement could be attributed to the physicochemical efficiency needed in disturbing the host membrane. Secondly, the electrostatic potential of the fusion peptide derived surface patches must be neutral or positive to be able to engage the host membrane. The presence of glycosylation sites adds to the hydrophobicity of the S2 domain surface, but it is a small fraction of the available surface for interaction with the host membrane. The fusogenic conformational transition requires optimal synergy between the physical and chemical properties of the fusion loop to allow a concurrent scything action to rapidly facilitate transition to the host-virus hemifusion membrane state. The free-energy available from the conformational transition of S2 to a more relaxed helical bundle is available to disrupt the host membrane and overcome the kinetic barrier to bring the host and virus membrane lipid bilayers together. The Contact Initiation Model ensures that the virus and host membrane are in close proximity for the formation of a hemifusion structure. The pre-hairpin S2 intermediate as suggested to exist by many researchers may be one of the many conformational states interacting with the host membrane. Priming by Spike cleavage is important for facilitating the rapid fusion process and therefore a part of the synergy at play. However, the open conformation of RBD seen in PDB ID: 6VSB for SARS-CoV-2 suggests that flexible linker segments loosely connect the RBD back to the fusion domain leaving it relatively free for unfettered conformational transition. Therefore, multifarious options to prime and trigger appear to be available to SARS-CoV-2 for viral entry, which contributes to its increased infectivity. Preventing the trigger by inhibiting the fusion loop is therefore a suitable target for therapy. Given the importance, a more extensive study of SARS-CoV-2 Spike protein and the mechanistic hypothesis described here is therefore warranted.

## Materials and Methods

The sequences used in this study were downloaded from the NCBI database (URL:http://www.ncbi.nlm.nih.gov). The multiple sequence alignments were performed using the T-coffee webserver (http://tcoffee.crg.cat/). The default parameter values for alignment available in the server were used. The server combines several methods to come up with an optimal multiple sequence alignment (42).

All protein three-dimensional structures were downloaded from the Protein Data Bank (http://www.rcsb.org). The PDB IDs for the downloaded structures are HCoV-HKU1: 5I08, MHV-A59: 3JCL, 6VSJ, HCoV-OC43: 6NZK, MERS-CoV: 6Q04, SARS-CoV-2: 6VXX, 6VSB, and SARS-CoV: 5XLR. Structures with the highest resolution was preferred when more than one model was available. Coordinates from these files were extracted for obtaining starting models of the FP-I, FP-II, FP-III, used in our molecular dynamics simulations. Whenever there were missing coordinates, they were modeled as an extended structure in the FP. The SASA was calculated by the program NACESS (http://wolf.bms.umist.ac.uk/naccess/); residues for which no atom coordinates are present in the PDB file was considered as fully exposed to solvent while calculating the relative SASA values. The secondary structure was calculated by the SECSTR program from the PROCHECK suite (https://www.ebi.ac.uk/thornton-srv/software/PROCHECK/). The electrostatic surface potential was calculated using the APBS plugin (38) inside the PyMol software (http://pymol.org). The default parameters were used for the calculations. All cartoon diagrams and surfaces were rendered by the PyMol software.

The molecular dynamics simulations were performed with GROMACS software (43) (http://www.gromacs.org). The simulations were performed using the CHARMM 27 forcefield with cmap (44). Each FP was placed in a cubic box solvated with water (SPC model). Solvent molecules were randomly replaced with Na^+^ and Cl^-^ to neutralize the system and bring the final concentration of NaCl to 0.1 M. Periodic boundary conditions were enforced in all three directions. The system was minimized until the maximum force in the system reached below 1000 kJ mol^-1^ nm^-1^. The systems were equilibrated for 2 ns under the NVT ensemble. This was followed by equilibration for 2 ns in the NPT ensemble at 1 atm and 300 K. The production run was executed for 500 ns saving the output every 100 ps yielding 5,000 frames for analysis. The analysis of the trajectory was performed using GROMACS utilities. The diagrams were created using our in-house software.

## Acknowledgments

DP thanks the Department of Biotechnology, New Delhi for supporting the computational facilities, and Prof. Jayasri Das Sarma, Department of Biological Sciences, Indian Institute of Science Education and Research Kolkata for her generous support.

## Supplementary Material

**Fig. S1.**
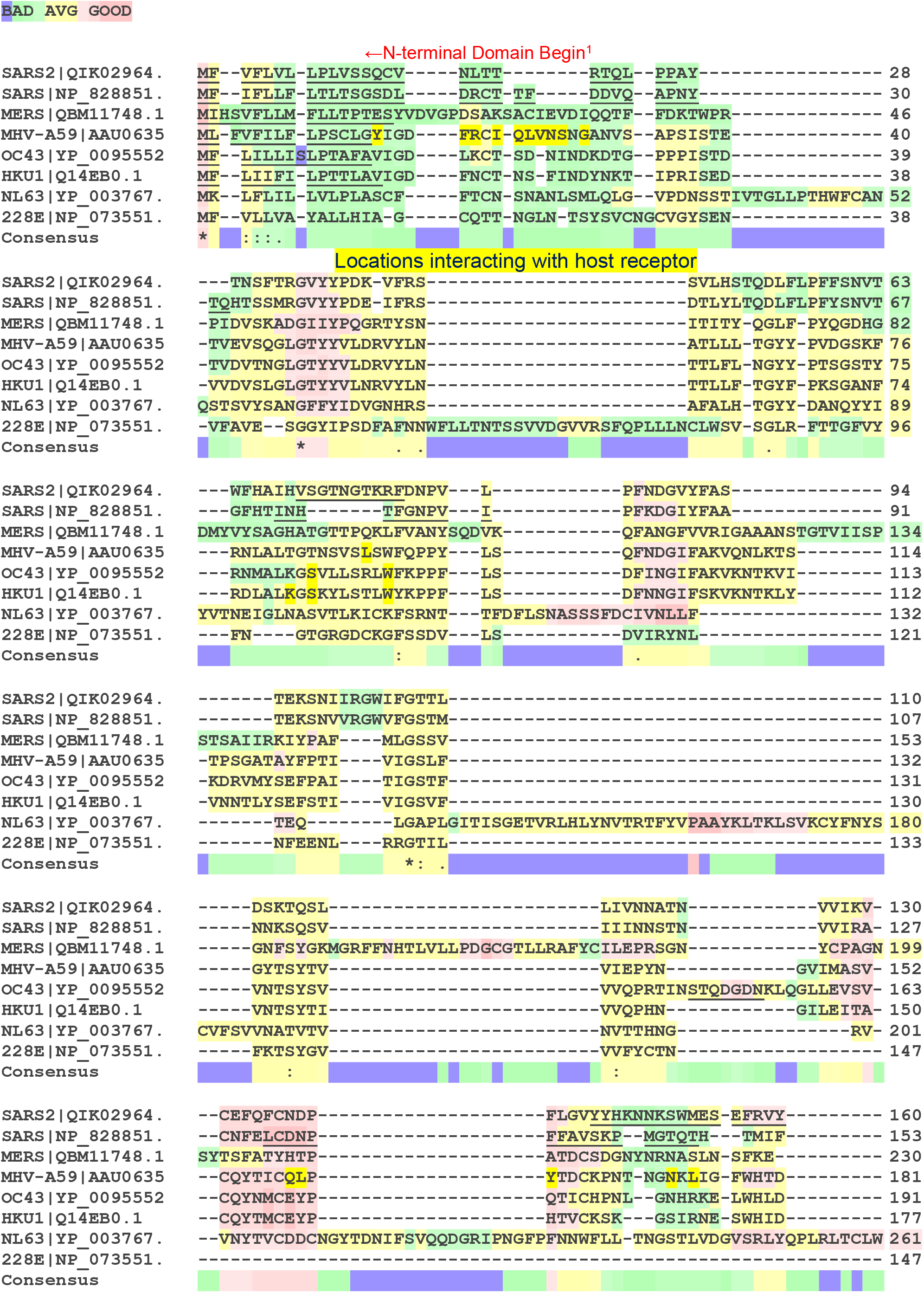
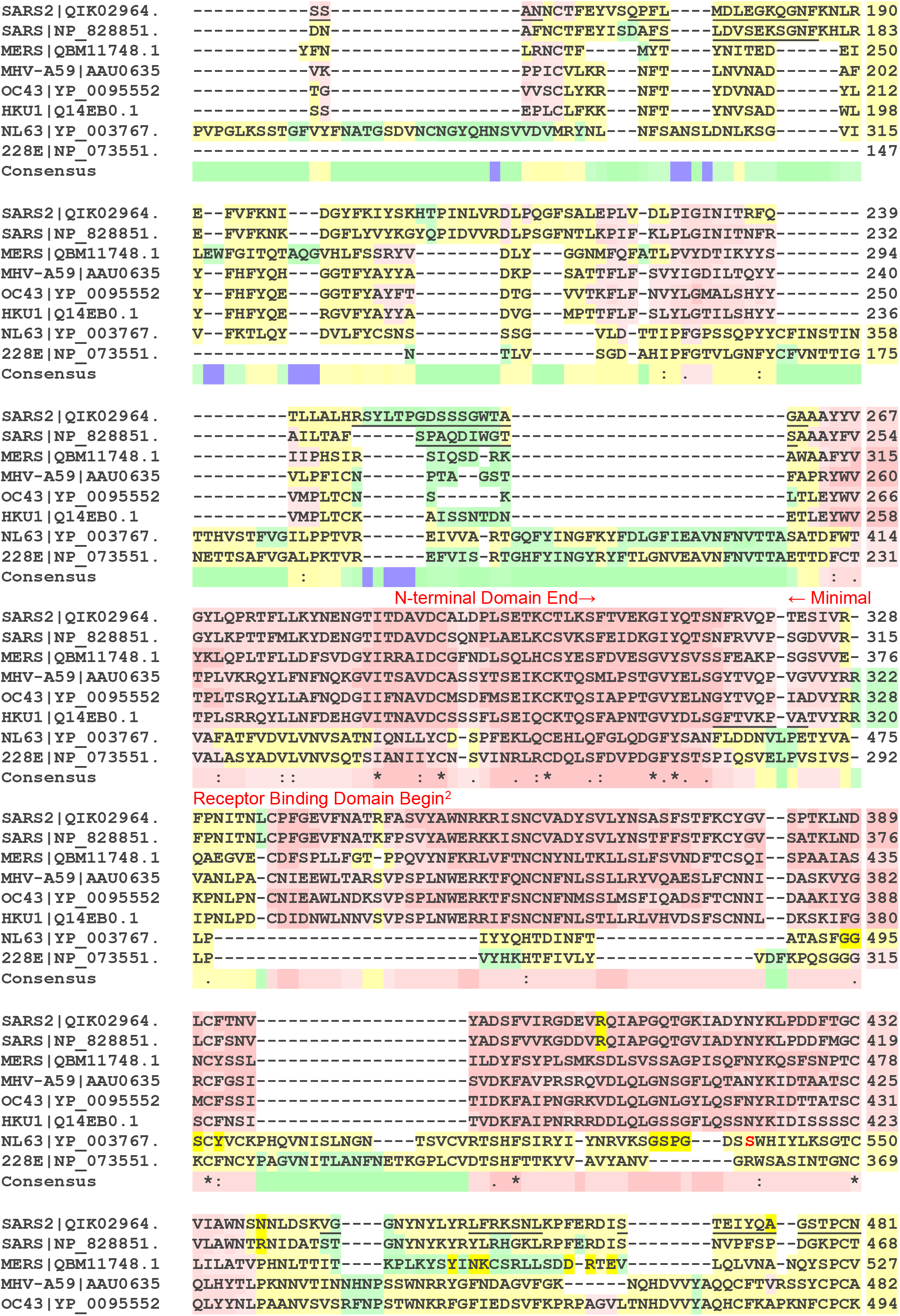
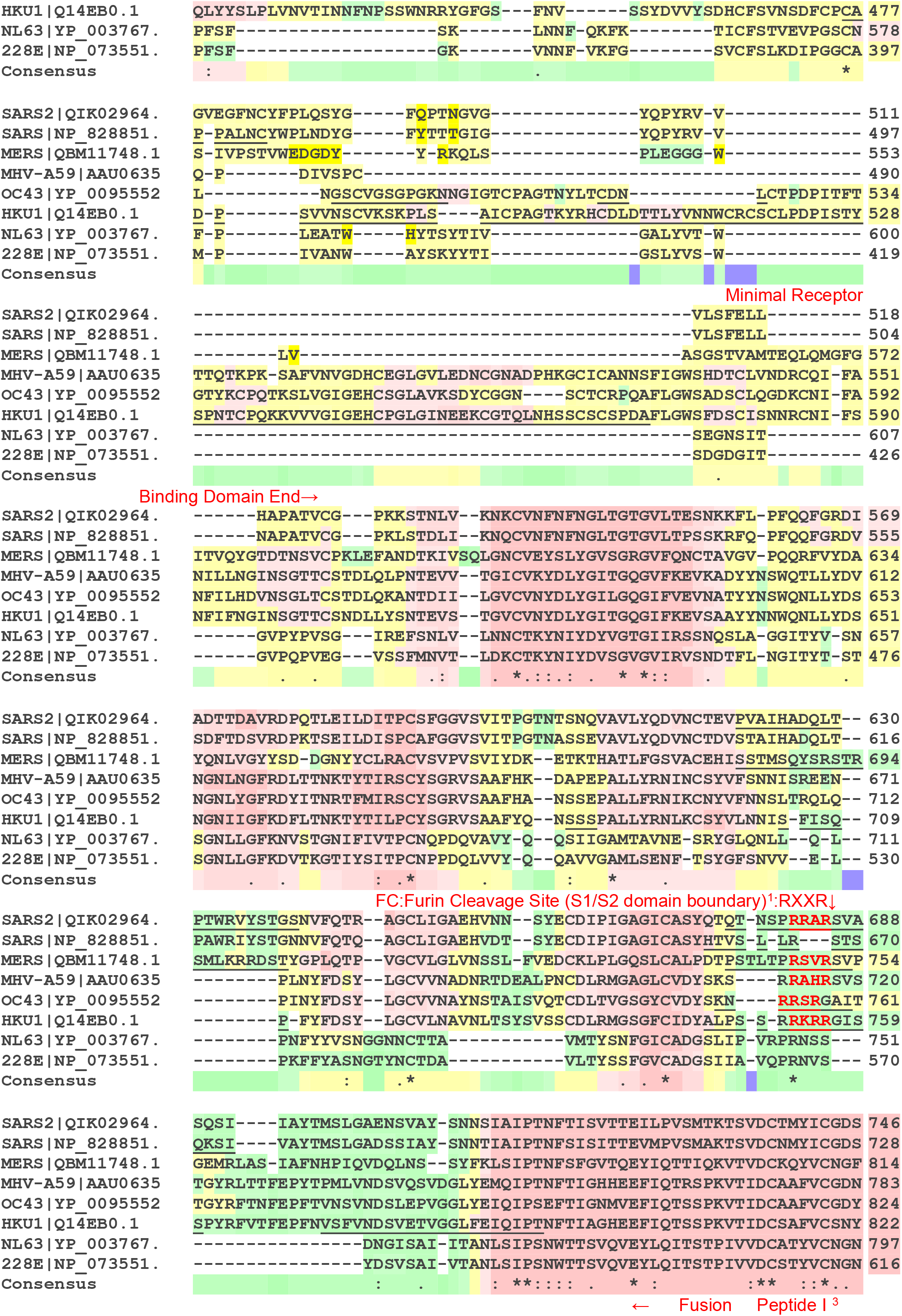
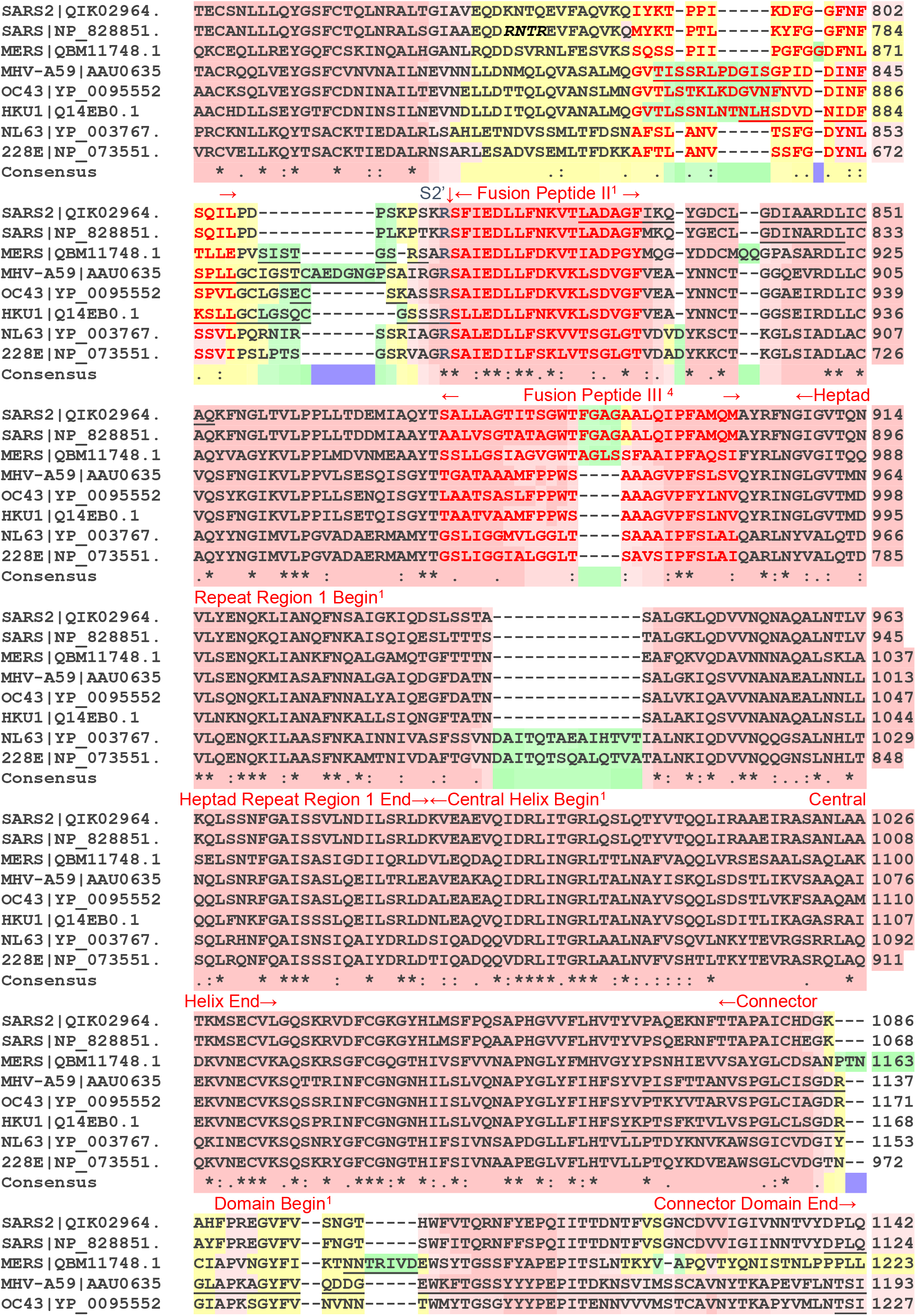
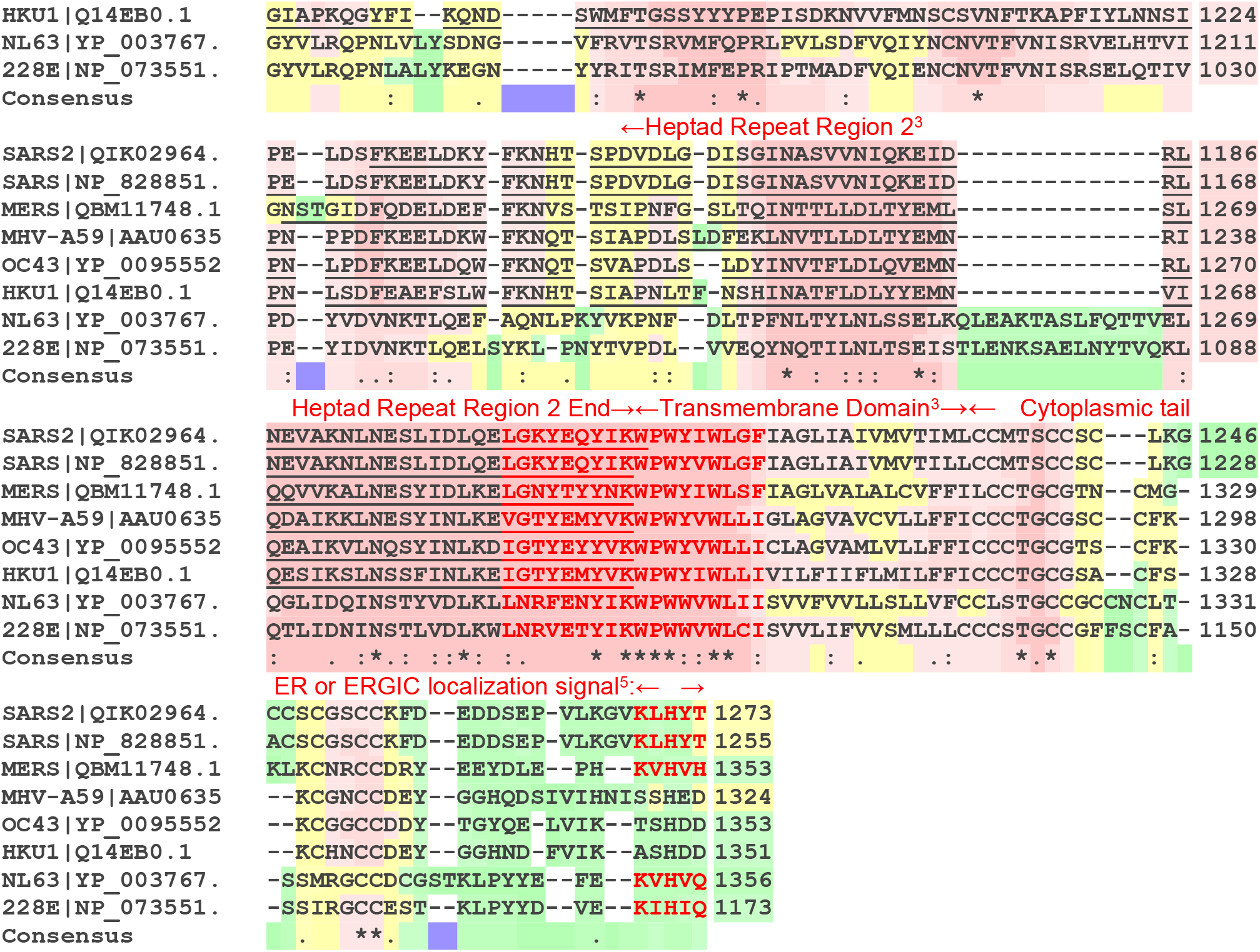
Multiple sequence alignment of Spike glycoproteins from eight coronaviruses. *Residues <u>underlined</u> have no coordinates in the structure file and are deemed flexible regions. The PDB files used to mark the same are SARS-CoV-2: 6VXX, SARS-CoV:5XLR, 6Q04: MERS-CoV, MHV-A59:3JCL, HCoV-OC43:6NZK, HCoV-HKU1:5I08. The sequences for HCoV-NL63 and HCoV-2289E are not marked because they belong to α-coronaviruses, whereas all others are from β-coronaviruses. The RED marked region spanning Heptad repeat region 2 C-terminal and trans-membrane domain N-terminal region is also called the aromatic domain or the fourth fusion peptide FP-IV*.

### Accession number of Genome/Spike protein sequences used in this alignment

1. AY700211.1 / AAU06356.1 (Murine hepatitis virus strain A59)
2. KT029139.1 / QBM11748.1 (MERS-CoV/KOR/KNIH/002_05_2015)
3. NC_004718.3 / NP_828851.1 (SARS coronavirus)
4. MN985325.1 / QIK02964.1 (2019-nCoV/USA-WA1/2020)
5. NC_005831.2 /YP_003767.1 (Human Coronavirus NL63,)
6. NC_006213.1 / YP_009555241.1 (Human coronavirus OC43 strain ATCC VR-759)
7. NC_006577.2 / Q14EB0.1 (Human coronavirus HKU1)
8. NC_002645.1 / NP_073551.1 (Human coronavirus 229E)

---

**Fig. S2.**
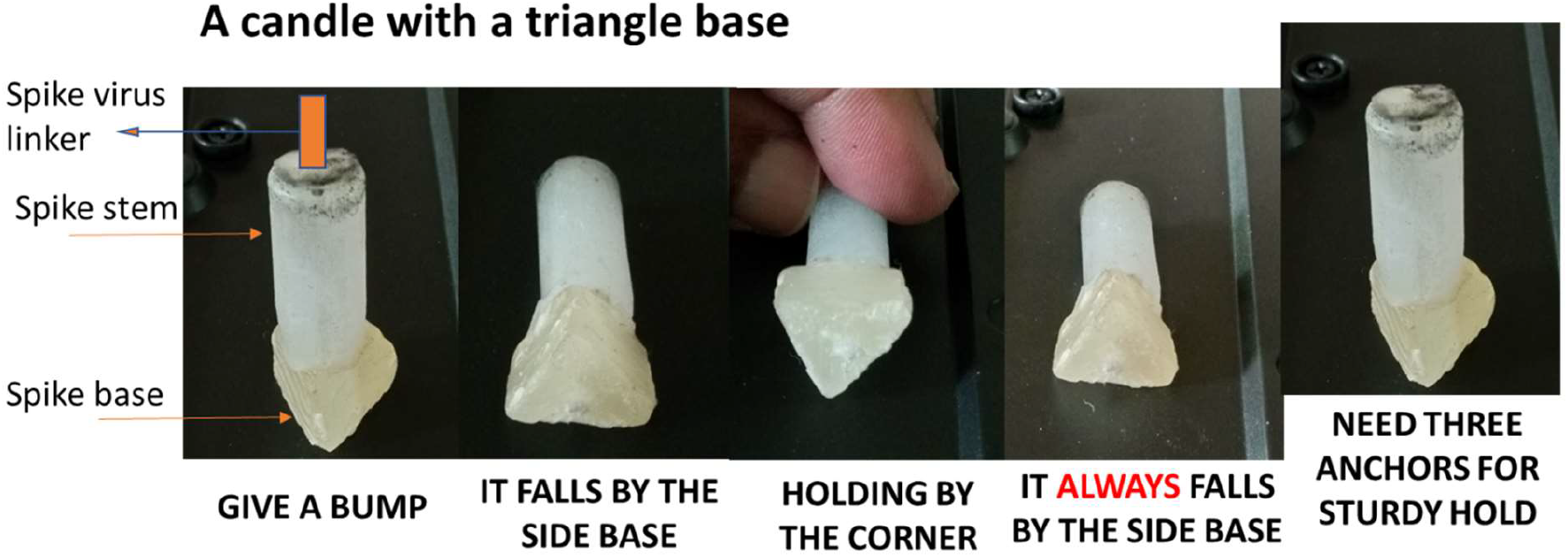
A series of snapshots showing the properties of the Contact Initiation Model. The model is created using candle wax, where the base represents the triangular shape as found for the trimeric spike S1 domain, and the cylindrical stem represents the trimeric S2 domain. The stem is attached to the virus envelope through trimeric flexible linkers at the C-terminal end of the Spike ectodomain. The model on the extreme left shows that an unanchored structure on a triangle base, when bumped from the side, will fall in a position where one of the base sides rests on the surface. This in turn would ensure that the fusion peptide surface on the stem aligned along the middle of the base will make the initial contact and trigger the fusion process (Please check the triangles marked in Fig. 2D alongside). The fact that the orientation holding it by the corner of the base is not stable is emphasized in third and fourth illustrations, and the model always falls such that it rests on the side base. This architecture guarantees the fusion peptides in the S2 domain make the initial contact given that they are located at the crest of the bulge on the S2 domain surface. The last figure emphasizes that three anchors are needed for a stable vertical “tripod” orientation of the spike with respect to the host surface.

**Fig. S3.**
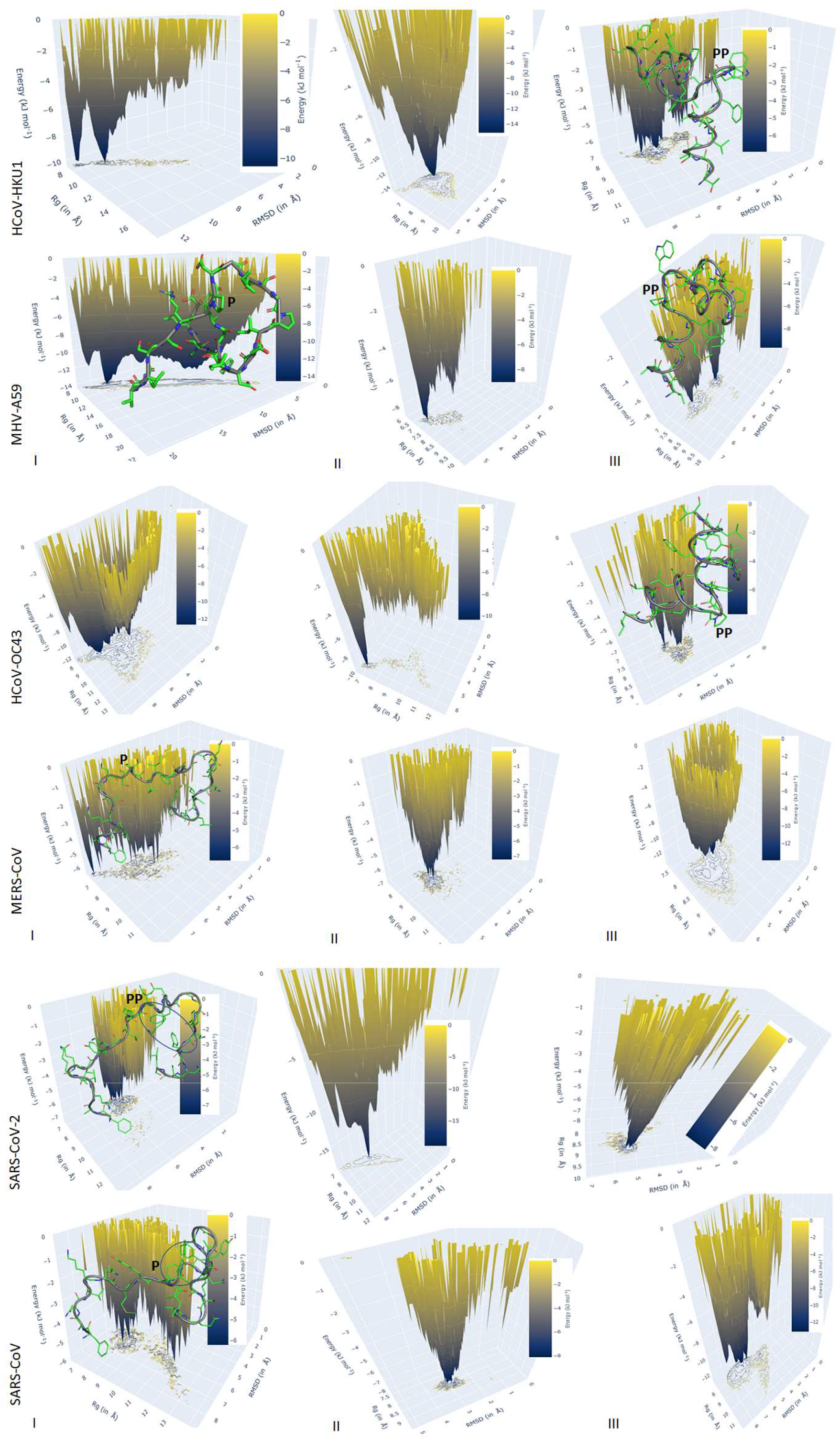
Energy landscape plot corresponding to all fusion peptides shown in Fig. 4 (main text). The ball and stick model of the starting structures of all the fusion peptides with central proline is shown along with their backbone trace. The central prolines are marked by “P”. Carbon atoms are in green, oxygen in red, nitrogen in blue, and sulfur in yellow. The location of the aromatic cluster is marked by oval in SARS-COV-2 and SARS-CoV Spike protein. The spread of the conformation in each FP can be visually estimated from the contours projected onto the X-Y plane.

