## Supplementary Figures for "Spike protein fusion loop controls SARS-CoV-2 fusogenicity and infectivity"

**Debnath Pal**

Department of Computational and Data Sciences, Indian Institute of Science, Bengaluru, Karnataka-560012, India

#### Supplementary Material

**Fig. S1**. **Multiple sequence alignment of Spike glycoproteins from eight coronaviruses.**

BAD AVG GOOD

```
                               ←N-terminal Domain Begin[1]
SARS2|QIK02964.    MF--VFLVL-LPLVSSQCV-----NLTT--------RTQL--PPAY----------------- 28
SARS|NP_828851.    MF--IFLLF-LTLTSGSDL-----DRCT-TF-----DDVQ--APNY----------------- 30
MERS|QBM11748.1    MIHSVFLLM-FLLTPTESYVDVGPDSAKSACIEVDIQQTF--FDKTWPR-------------- 46
MHV-A59|AAU0635    ML-FVFILF-LPSCLGYIGD----FRCI-QLVNSNGANVS--APSISTE-------------- 40
OC43|YP_0095552    MF--LILLISLPTAFAVIGD----LKCT-SD-NINDKDTG--PPPISTD-------------- 39
HKU1|Q14EB0.1      MF--LIIFI-LPTTLAVIGD----FNCT-NS-FINDYNKT--IPRISED-------------- 38
NL63|YP_003767.    MK--LFLIL-LVLPLASCF-----FTCN-SNANLSMLQLG--VPDNSSTIVTGLLPTHWFCAN 52
228E|NP_073551.    MF--VLLVA-YALLHIA-G-----CQTT-NGLN-TSYSVCNGCVGYSEN-------------- 38
Consensus          *   :::.
                         Locations interacting with host receptor
SARS2|QIK02964.    ---TNSFTRGVYYPDK-VFRS------------------------SVLHSTQDLFLPFFSNVT 63
SARS|NP_828851.    -TQHTSSMRGVYYPDE-IFRS------------------------DTLYLTQDLFLPFYSNVT 67
MERS|QBM11748.1    -PIDVSKADGIIYPQGRTYSN------------------------ITITY-QGLF-PYQGDHG 82
MHV-A59|AAU0635    -TVEVSQGLGTYYVLDRVYLN------------------------ATLLL-TGYY-PVDGSKF 76
OC43|YP_0095552    -TVDVTNGLGTYYVLDRVYLN------------------------TTLFL-NGYY-PTSGSTY 75
HKU1|Q14EB0.1      -VVDVSLGLGTYYVLNRVYLN------------------------TTLLF-TGYF-PKSGANF 74
NL63|YP_003767.    QSTSVYSANGFFYIDVGNHRS------------------------AFALH-TGYY-DANQYYI 89
228E|NP_073551.    -VFAVE--SGGYIPSDFAFNNWFLLTNTSSVVDGVVRSFQPLLLNCLWSV-SGLR-FTTGFVY 96
Consensus                   *        . .                               .

SARS2|QIK02964.    ---WFHAIHVSGTNGTKRFDNPV---L------------PFNDGVYFAS-------------- 94
SARS|NP_828851.    ---GFHTINH-------TFGNPV---I------------PFKDGIYFAA-------------- 91
MERS|QBM11748.1    DMYVYSAGHATGTTPQKLFVANYSQDVK-----------QFANGFVVRIGAAANSTGTVIISP 134
MHV-A59|AAU0635    ---RNLALTGTNSVSLSWFQPPY---LS-----------QFNDGIFAKVQNLKTS-------- 114
OC43|YP_0095552    ---RNMALKGSVLLSRLWFKPPF---LS-----------DFINGIFAKVKNTKVI-------- 113
HKU1|Q14EB0.1      ---RDLALKGSKYLSTLWYKPPF---LS-----------DFNNGIFSKVKNTKLY-------- 112
NL63|YP_003767.    YVTNEIGLNASVTLKICKFSRNT---TFDFLSNASSSFDCIVNLLF----------------- 132
228E|NP_073551.    ---FN----GTGRGDCKGFSSDV---LS-----------DVIRYNL----------------- 121
Consensus                            :                     .

SARS2|QIK02964.    -------TEKSNIIRGWIFGTTL---------------------------------------- 110
SARS|NP_828851.    -------TEKSNVVRGWVFGSTM---------------------------------------- 107
MERS|QBM11748.1    STSAIIRKIYPAF----MLGSSV---------------------------------------- 153
MHV-A59|AAU0635    -TPSGATAYFPTI----VIGSLF---------------------------------------- 132
OC43|YP_0095552    -KDRVMYSEFPAI----TIGSTF---------------------------------------- 131
HKU1|Q14EB0.1      -VNNTLYSEFSTI----VIGSVF---------------------------------------- 130
NL63|YP_003767.    -------TEQ--------LGAPLGITISGETVRLHLYNVTRTFYVPAAYKLTKLSVKCYFNYS 180
228E|NP_073551.    -------NFEENL----RRGTIL---------------------------------------- 133
Consensus                             *: .

SARS2|QIK02964.    -----DSKTQSL-------------------------LIVNNATN------------VVIKV- 130
SARS|NP_828851.    -----NNKSQSV-------------------------IIINNSTN------------VVIRA- 127
MERS|QBM11748.1    -----GNFSYGKMGRFFNHTLVLLPDGCGTLLRAFYCILEPRSGN------------YCPAGN 199
MHV-A59|AAU0635    -----GYTSYTV-------------------------VIEPYN------------GVIMASV- 152
OC43|YP_0095552    -----VNTSYSV-------------------------VVQPRTINSTQDGDNKLQGLLEVSV- 163
HKU1|Q14EB0.1      -----VNTSYTI-------------------------VVQPHN------------GILEITA- 150
NL63|YP_003767.    CVFSVVNATVTV-------------------------NVTTHNG----------------RV- 201
228E|NP_073551.    -----FKTSYGV-------------------------VVFYCTN------------------- 147
Consensus                  :                             :

SARS2|QIK02964.    --CEFQFCNDP---------------------FLGVYYHKNNKSWMES-EFRVY--------- 160
SARS|NP_828851.    --CNFELCDNP---------------------FFAVSKP--MGTQTH---TMIF--------- 153
MERS|QBM11748.1    SYTSFATYHTP---------------------ATDCSDGNYNRNASLN-SFKE---------- 230
MHV-A59|AAU0635    --CQYTICQLP---------------------YTDCKPNT-NGNKLIG-FWHTD--------- 181
OC43|YP_0095552    --CQYNMCEYP---------------------QTICHPNL--GNHRKE-LWHLD--------- 191
HKU1|Q14EB0.1      --CQYTMCEYP---------------------HTVCKSK---GSIRNE-SWHID--------- 177
NL63|YP_003767.    --VNYTVCDDCNGYTDNIFSVQQDGRIPNGFPFNNWFLL-TNGSTLVDGVSRLYQPLRLTCLW 261
228E|NP_073551.    --------------------------------------------------------------- 147
Consensus
```

### Supplementary Material

```
SARS2|QIK02964.   -----------SS------------------ANNCTFEYVSQPFL----MDLEGKQGNFKNLR 190
SARS|NP_828851.   -----------DN------------------AFNCTFEYISDAFS----LDVSEKSGNFKHLR 183
MERS|QBM11748.1   ----------YFN------------------LRNCTF-----MYT----YNITED------EI 250
MHV-A59|AAU0635   -----------VK------------------PPICVLKR---NFT----LNVNAD------AF 202
OC43|YP_0095552   -----------TG------------------VVSCLYKR---NFT----YDVNAD------YL 212
HKU1|Q14EB0.1     -----------SS------------------EPLCLFKK---NFT----YNVSAD------WL 198
NL63|YP_003767.   PVPGLKSSTGFVYFNATGSDVNCNGYQHNSVVDVMRYNL---NFSANSLDNLKSG------VI 315
228E|NP_073551.   --------------------------------------------------------------- 147
Consensus

SARS2|QIK02964.   E--FVFKNI---DGYFKIYSKHTPINLVRDLPQGFSALEPLV-DLPIGINITRFQ-------- 239
SARS|NP_828851.   E--FVFKNK---DGFLYVYKGYQPIDVVRDLPSGFNTLKPIF-KLPLGINITNFR-------- 232
MERS|QBM11748.1   LEWFGITQTAQGVHLFSSRYV--------DLY---GGNMFQFATLPVYDTIKYYS-------- 294
MHV-A59|AAU0635   Y--FHFYQH---GGTFYAYYA--------DKP---SATTFLF-SVYIGDILTQYY-------- 240
OC43|YP_0095552   Y--FHFYQE---GGTFYAYFT--------DTG---VVTKFLF-NVYLGMALSHYY-------- 250
HKU1|Q14EB0.1     Y--FHFYQE---RGVFYAYYA--------DVG---MPTTFLF-SLYLGTILSHYY-------- 236
NL63|YP_003767.   V--FKTLQY---DVLFYCSNS--------SSG------VLD-TTIPFGPSSQPYYCFINSTIN 358
228E|NP_073551.   --------------------N--------TLV------SGD-AHIPFGTVLGNFYCFVNTTIG 175
Consensus                                                     : .      :

SARS2|QIK02964.   ---------TLLALHRSYLTPGDSSSGWTA--------------------------GAAAYYV 267
SARS|NP_828851.   ---------AILTAF------SPAQDIWGT--------------------------SAAAYFV 254
MERS|QBM11748.1   ---------IIPHSIR------SIQSD-RK--------------------------AWAAFYV 315
MHV-A59|AAU0635   ---------VLPFICN------PTA--GST--------------------------FAPRYWV 260
OC43|YP_0095552   ---------VMPLTCN------S------K--------------------------LTLEYWV 266
HKU1|Q14EB0.1     ---------VMPLTCK------AISSNTDN--------------------------ETLEYWV 258
NL63|YP_003767.   TTHVSTFVGILPPTVR------EIVVA-RTGQFYINGFKYFDLGFIEAVNFNVTTASATDFWT 414
228E|NP_073551.   NETTSAFVGALPKTVR------EFVIS-RTGHFYINGYRYFTLGNVEAVNFNVTTAETTDFCT 231
Consensus                   :                                                 : .
```

N-terminal Domain End→ ← Minimal

```
SARS2|QIK02964.   GYLQPRTFLLKYNENGTITDAVDCALDPLSETKCTLKSFTVEKGIYQTSNFRVQP-TESIVR- 328
SARS|NP_828851.   GYLKPTTFMLKYDENGTITDAVDCSQNPLAELKCSVKSFEIDKGIYQTSNFRVVP-SGDVVR- 315
MERS|QBM11748.1   YKLQPLTFLLDFSVDGYIRRAIDCGFNDLSQLHCSYESFDVESGVYSVSSFEAKP-SGSVVE- 376
MHV-A59|AAU0635   TPLVKRQYLFNFNQKGVITSAVDCASSYTSEIKCKTQSMLPSTGVYELSGYTVQP-VGVVYRR 322
OC43|YP_0095552   TPLTSRQYLLAFNQDGIIFNAVDCMSDFMSEIKCKTQSIAPPTGVYELNGYTVQP-IADVYRR 328
HKU1|Q14EB0.1     TPLSRRQYLLNFDEHGVITNAVDCSSSFLSEIQCKTQSFAPNTGVYDLSGFTVKP-VATVYRR 320
NL63|YP_003767.   VAFATFVDVLVNVSATNIQNLLYCD-SPFEKLQCEHLQFGLQDGFYSANFLDDNVLPETYVA- 475
228E|NP_073551.   VALASYADVLVNVSQTSIANIIYCN-SVINRLRCDQLSFDVPDGFYSTSPIQSVELPVSIVS- 292
Consensus           :     ::       *   : *  .   . :*   .:    *.*. .
```

Receptor Binding Domain Begin[2]

```
SARS2|QIK02964.   FPNITNLCPFGEVFNATRFASVYAWNRKRISNCVADYSVLYNSASFSTFKCYGV--SPTKLND 389
SARS|NP_828851.   FPNITNLCPFGEVFNATKFPSVYAWERKKISNCVADYSVLYNSTFFSTFKCYGV--SATKLND 376
MERS|QBM11748.1   QAEGVE-CDFSPLLFGT-PPQVYNFKRLVFTNCNYNLTKLLSLFSVNDFTCSQI--SPAAIAS 435
MHV-A59|AAU0635   VANLPA-CNIEEWLTARSVPSPLNWERKTFQNCNFNLSSLLRYVQAESLFCNNI--DASKVYG 382
OC43|YP_0095552   KPNLPN-CNIEAWLNDKSVPSPLNWERKTFSNCNFNMSSLMSFIQADSFTCNNI--DAAKIYG 388
HKU1|Q14EB0.1     IPNLPD-CDIDNWLNNVSVPSPLNWERRIFSNCNFNLSTLLRLVHVDSFSCNNL--DKSKIFG 380
NL63|YP_003767.   LP-------------------------IYYQHTDINFT------------------ATASFGG 495
228E|NP_073551.   LP-------------------------VYHKHTFIVLY---------------VDFKPQSGGG 315
Consensus          .                             :                              .

SARS2|QIK02964.   LCFTNV--------------------YADSFVIRGDEVRQIAPGQTGKIADYNYKLPDDFTGC 432
SARS|NP_828851.   LCFSNV--------------------YADSFVVKGDDVRQIAPGQTGVIADYNYKLPDDFMGC 419
MERS|QBM11748.1   NCYSSL--------------------ILDYFSYPLSMKSDLSVSSAGPISQFNYKQSFSNPTC 478
MHV-A59|AAU0635   RCFGSI--------------------SVDKFAVPRSRQVDLQLGNSGFLQTANYKIDTAATSC 425
OC43|YP_0095552   MCFSSI--------------------TIDKFAIPNGRKVDLQLGNLGYLQSFNYRIDTTATSC 431
HKU1|Q14EB0.1     SCFNSI--------------------TVDKFAIPNRRRDDLQLGSSGFLQSSNYKIDISSSSC 423
NL63|YP_003767.   SCYVCKPHQVNISLNGN----TSVCVRTSHFSIRYI-YNRVKSGSPG---DSSWHIYLKSGTC 550
228E|NP_073551.   KCFNCYPAGVNITLANFNETKGPLCVDTSHFTTKYV-AVYANV--------GRWSASINTGNC 369
Consensus          *:                         . *                      :        *

SARS2|QIK02964.   VIAWNSNNLDSKVG----GNYNYLYRLFRKSNLKPFERDIS--------TEIYQA--GSTPCN 481
SARS|NP_828851.   VLAWNTRNIDATST----GNYNYKYRYLRHGKLRPFERDIS--------NVPFSP--DGKPCT 468
MERS|QBM11748.1   LILATVPHNLTTIT----KPLKYSYINKCSRLLSDD-RTEV--------LQLVNA-NQYSPCV 527
MHV-A59|AAU0635   QLHYTLPKNNVTINNHNPSSWNRRYGFNDAGVFGK------NQHDVVYAQQCFTVRSSYCPCA 482
OC43|YP_0095552   QLYYNLPAANVSVSRFNPSTWNKRFGFIEDSVFKPRPAGVLTNHDVVYAQHCFKAPKNFCPCK 494
```

#### Supplementary Material

```
HKU1|Q14EB0.1      QLYYSLPLVNVTINNFNPSSWNRRYGFGS---FNV------SSYDVVYSDHCFSVNSDFCPCA 477
NL63|YP_003767.    PFSF-------------------SK-------LNNF-QKFK--------TICFSTVEVPGSCN 578
228E|NP_073551.    PFSF-------------------GK-------VNNF-VKFG--------SVCFSLKDIPGGCA 397
Consensus           :                              .                            *

SARS2|QIK02964.    GVEGFNCYFPLQSYG-----FQPTNGVG--------------YQPYRV-V------------- 511
SARS|NP_828851.    P-PALNCYWPLNDYG-----FYTTTGIG--------------YQPYRV-V------------- 497
MERS|QBM11748.1    S-IVPSTVWEDGDY-------Y-RKQLS--------------PLEGGG-W------------- 553
MHV-A59|AAU0635    Q-P-------DIVSPC----------------------------------------------- 490
OC43|YP_0095552    L-----------NGSCVGSGPGKNNGIGTCPAGTNYLTCDN------------LCTPDPITFT 534
HKU1|Q14EB0.1      D-P-------SVVNSCVKSKPLS----AICPAGTKYRHCDLDTTLYVNNWCRCSCLPDPISTY 528
NL63|YP_003767.    F-P-------LEATW-----HYTSYTIV--------------GALYVT-W------------- 600
228E|NP_073551.    M-P-------IVANW-----AYSKYYTI--------------GSLYVS-W------------- 419
Consensus
                                                             Minimal Receptor
SARS2|QIK02964.    -----------------------------------------------VLSFELL--------- 518
SARS|NP_828851.    -----------------------------------------------VLSFELL--------- 504
MERS|QBM11748.1    --------LV------------------------------------ASGSTVAMTEQLQMGFG 572
MHV-A59|AAU0635    TTQTKPK-SAFVNVGDHCEGLGVLEDNCGNADPHKGCICANNSFIGWSHDTCLVNDRCQI-FA 551
OC43|YP_0095552    GTYKCPQTKSLVGIGEHCSGLAVKSDYCGGN----SCTCRPQAFLGWSADSCLQGDKCNI-FA 592
HKU1|Q14EB0.1      SPNTCPQKKVVVGIGEHCPGLGINEEKCGTQLNHSSCSCSPDAFLGWSFDSCISNNRCNI-FS 590
NL63|YP_003767.    -----------------------------------------------SEGNSIT--------- 607
228E|NP_073551.    -----------------------------------------------SDGDGIT--------- 426
Consensus                                                           .
               Binding Domain End→
SARS2|QIK02964.    ------HAPATVCG---PKKSTNLV--KNKCVNFNFNGLTGTGVLTESNKKFL-PFQQFGRDI 569
SARS|NP_828851.    ------NAPATVCG---PKLSTDLI--KNQCVNFNFNGLTGTGVLTPSSKRFQ-PFQQFGRDV 555
MERS|QBM11748.1    ITVQYGTDTNSVCPKLEFANDTKIVSQLGNCVEYSLYGVSGRGVFQNCTAVGV-PQQRFVYDA 634
MHV-A59|AAU0635    NILLNGINSGTTCSTDLQLPNTEVV--TGICVKYDLYGITGQGVFKEVKADYYNSWQTLLYDV 612
OC43|YP_0095552    NFILHDVNSGLTCSTDLQKANTDII--LGVCVNYDLYGILGQGIFVEVNATYYNSWQNLLYDS 653
HKU1|Q14EB0.1      NFIFNGINSGTTCSNDLLYSNTEVS--TGVCVNYDLYGITGQGIFKEVSAAYYNNWQNLLYDS 651
NL63|YP_003767.    ------GVPYPVSG---IREFSNLV--LNNCTKYNIYDYVGTGIIRSSNQSLA-GGITYV-SN 657
228E|NP_073551.    ------GVPQPVEG---VSSFMNVT--LDKCTKYNIYDVSGVGVIRVSNDTFL-NGITYT-ST 476
Consensus                  .  .          .:    . *.::.: .  * *::   .            .

SARS2|QIK02964.    ADTTDAVRDPQTLEILDITPCSFGGVSVITPGTNTSNQVAVLYQDVNCTEVPVAIHADQLT-- 630
SARS|NP_828851.    SDFTDSVRDPKTSEILDISPCAFGGVSVITPGTNASSEVAVLYQDVNCTDVSTAIHADQLT-- 616
MERS|QBM11748.1    YQNLVGYYSD-DGNYYCLRACVSVPVSVIYDK--ETKTHATLFGSVACEHISSTMSQYSRSTR 694
MHV-A59|AAU0635    NGNLNGFRDLTTNKTYTIRSCYSGRVSAAFHK--DAPEPALLYRNINCSYVFSNNISREEN-- 671
OC43|YP_0095552    NGNLYGFRDYITNRTFMIRSCYSGRVSAAFHA--NSSEPALLFRNIKCNYVFNNSLTRQLQ-- 712
HKU1|Q14EB0.1      NGNIIGFKDFLTNKTYTILPCYSGRVSAAFYQ--NSSSPALLYRNLKCSYVLNNIS-FISQ-- 709
NL63|YP_003767.    SGNLLGFKNVSTGNIFIVTPCNQPDQVAVY-Q--QSIIGAMTAVNE-SRYGLQNLL--Q-L-- 711
228E|NP_073551.    SGNLLGFKDVTKGTIYSITPCNPPDQLVVY-Q--QAVVGAMLSENF-TSYGFSNVV--E-L-- 530
Consensus               .  .        : .*      .       :   *    .
                                  FC:Furin Cleavage Site (S1/S2 domain boundary)[1]:RXXR↓
SARS2|QIK02964.    PTWRVYSTGSNVFQTR--AGCLIGAEHVNN--SYECDIPIGAGICASYQTQT-NSPRRARSVA 688
SARS|NP_828851.    PAWRIYSTGNNVFQTQ--AGCLIGAEHVDT--SYECDIPIGAGICASYHTVS-L-LR---STS 670
MERS|QBM11748.1    SMLKRRDSTYGPLQTP--VGCVLGLVNSSL-FVEDCKLPLGQSLCALPDTPSTLTPRSVRSVP 754
MHV-A59|AAU0635    --------PLNYFDSY--LGCVVNADNRTDEALPNCDLRMGAGLCVDYSKS----RRAHRSVS 720
OC43|YP_0095552    --------PINYFDSY--LGCVVNAYNSTAISVQTCDLTVGSGYCVDYSKN----RRSRGAIT 761
HKU1|Q14EB0.1      --------P-FYFDSY--LGCVLNAVNLTSYSVSSCDLRMGSGFCIDYALPS-S-RRKRRGIS 759
NL63|YP_003767.    --------PNFYYVSNGGNNCTTA-----------VMTYSNFGICADGSLIP-VRPRNSS--- 751
228E|NP_073551.    --------PKFFYASNGTYNCTDA-----------VLTYSSFGVCADGSIIA-VQPRNVS--- 570
Consensus                        :    .*                   . . *           *

SARS2|QIK02964.    SQSI----IAYTMSLGAENSVAY-SNNSIAIPTNFTISVTTEILPVSMTKTSVDCTMYICGDS 746
SARS|NP_828851.    QKSI----VAYTMSLGADSSIAY-SNNTIAIPTNFSISITTEVMPVSMAKTSVDCNMYICGDS 728
MERS|QBM11748.1    GEMRLAS-IAFNHPIQVDQLNS--SYFKLSIPTNFSFGVTQEYIQTTIQKVTVDCKQYVCNGF 814
MHV-A59|AAU0635    TGYRLTTFEPYTPMLVNDSVQSVDGLYEMQIPTNFTIGHHEEFIQTRSPKVTIDCAAFVCGDN 783
OC43|YP_0095552    TGYRFTNFEPFTVNSVNDSLEPVGGLYEIQIPSEFTIGNMVEFIQTSSPKVTIDCAAFVCGDY 824
HKU1|Q14EB0.1      SPYRFVTFEPFNVSFVNDSVETVGGLFEIQIPTNFTIAGHEEFIQTSSPKVTIDCSAFVCSNY 822
NL63|YP_003767.    ----------------DNGISAI-ITANLSIPSNWTTSVQVEYLQITSTPIVVDCATYVCNGN 797
228E|NP_073551.    ----------------YDSVSAI-VTANLSIPSNWTTSVQVEYLQITSTPIVVDCSTYVCNGN 616
Consensus                           :   .      : **:::: .   * :        :**  ::*..
                                                    ←      Fusion     Peptide I [3]
```

### Supplementary Material

```
SARS2|QIK02964.   TECSNLLLQYGSFCTQLNRALTGIAVEQDKNTQEVFAQVKQIYKT-PPI-----KDFG-GFNF 802
SARS|NP_828851.   TECANLLLQYGSFCTQLNRALSGIAAEQDRNTREVFAQVKQMYKT-PTL-----KYFG-GFNF 784
MERS|QBM11748.1   QKCEQLLREYGQFCSKINQALHGANLRQDDSVRNLFESVKSSQSS-PII-----PGFGGDFNL 871
MHV-A59|AAU0635   TACRQQLVEYGSFCVNVNAILNEVNNLLDNMQLQVASALMQGVTISSRLPDGISGPID-DINF 845
OC43|YP_0095552   AACKSQLVEYGSFCDNINAILTEVNELLDTTQLQVANSLMNGVTLSTKLKDGVNFNVD-DINF 886
HKU1|Q14EB0.1     AACHDLLSEYGTFCDNINSILNEVNDLLDITQLQVANALMQGVTLSSNLNTNLHSDVD-NIDF 884
NL63|YP_003767.   PRCKNLLKQYTSACKTIEDALRLSAHLETNDVSSMLTFDSNAFSL-ANV-----TSFG-DYNL 853
228E|NP_073551.   VRCVELLKQYTSACKTIEDALRNSARLESADVSEMLTFDKKAFTL-ANV-----SSFG-DYNL 672
Consensus          * . * :*   *  ::  *              .:       .  . .  :     ..  . ::

                   →                    S2'↓← Fusion Peptide II[1] →
SARS2|QIK02964.   SQILPD-----------PSKPSKRSFIEDLLFNKVTLADAGFIKQ-YGDCL--GDIAARDLIC 851
SARS|NP_828851.   SQILPD-----------PLKPTKRSFIEDLLFNKVTLADAGFMKQ-YGECL--GDINARDLIC 833
MERS|QBM11748.1   TLLEPVSIST-------GS-RSARSAIEDLLFDKVTIADPGYMQG-YDDCMQQGPASARDLIC 925
MHV-A59|AAU0635   SPLLGCIGSTCAEDGNGPSAIRGRSAIEDLLFDKVKLSDVGFVEA-YNNCT--GGQEVRDLLC 905
OC43|YP_0095552   SPVLGCLGSEC-------SKASSRSAIEDLLFDKVKLSDVGFVEA-YNNCT--GGAEIRDLIC 939
HKU1|Q14EB0.1     KSLLGCLGSQC--------GSSSRSLLEDLLFNKVKLSDVGFVEA-YNNCT--GGSEIRDLLC 936
NL63|YP_003767.   SSVLPQRNIR-------SSRIAGRSALEDLLFSKVVTSGLGTVDVDYKSCT--KGLSIADLAC 907
228E|NP_073551.   SSVIPSLPTS-------GSRVAGRSAIEDILFSKLVTSGLGTVDADYKKCT--KGLSIADLAC 726
Consensus         . :                      ** :**:**.*:    :.  *  :.    * .*        ** *

                                        ←        Fusion Peptide III [4]       →     ←Heptad
SARS2|QIK02964.   AQKFNGLTVLPPLLTDEMIAQYTSALLAGTITSGWTFGAGAALQIPFAMQMAYRFNGIGVTQN 914
SARS|NP_828851.   AQKFNGLTVLPPLLTDDMIAAYTAALVSGTATAGWTFGAGAALQIPFAMQMAYRFNGIGVTQN 896
MERS|QBM11748.1   AQYVAGYKVLPPLMDVNMEAAYTSSLLGSIAGVGWTAGLSSFAAIPFAQSIFYRLNGVGITQQ 988
MHV-A59|AAU0635   VQSFNGIKVLPPVLSESQISGYTTGATAAAMFPPWS----AAAGVPFSLSVQYRINGLGVTMN 964
OC43|YP_0095552   VQSYKGIKVLPPLLSENQISGYTLAATSASLFPPWT----AAAGVPFYLNVQYRINGLGVTMD 998
HKU1|Q14EB0.1     VQSFNGIKVLPPILSETQISGYTTAATVAAMFPPWS----AAAGVPFSLNVQYRINGLGVTMD 995
NL63|YP_003767.   AQYYNGIMVLPGVADAERMAMYTGSLIGGMVLGGLT----SAAAIPFSLALQARLNYVALQTD 966
228E|NP_073551.   AQYYNGIMVLPGVADAERMAMYTGSLIGGIALGGLT----SAVSIPFSLAIQARLNYVALQTD 785
Consensus         .*    *   ***         :  **  .    .        :     :  :**    :    *:*  :.:   :

                  Repeat Region 1 Begin[1]
SARS2|QIK02964.   VLYENQKLIANQFNSAIGKIQDSLSSTA--------------SALGKLQDVVNQNAQALNTLV 963
SARS|NP_828851.   VLYENQKQIANQFNKAISQIQESLTTTS--------------TALGKLQDVVNQNAQALNTLV 945
MERS|QBM11748.1   VLSENQKLIANKFNQALGAMQTGFTTTN--------------EAFQKVQDAVNNNAQALSKLA 1037
MHV-A59|AAU0635   VLSENQKMIASAFNNALGAIQDGFDATN--------------SALGKIQSVVNANAEALNNLL 1013
OC43|YP_0095552   VLSQNQKLIANAFNNALYAIQEGFDATN--------------SALVKIQAVVNANAEALNNLL 1047
HKU1|Q14EB0.1     VLNKNQKLIANAFNKALLSIQNGFTATN--------------SALAKIQSVVNANAQALNSLL 1044
NL63|YP_003767.   VLQENQKILAASFNKAINNIVASFSSVNDAITQTAEAIHTVTIALNKIQDVVNQQGSALNHLT 1029
228E|NP_073551.   VLQENQKILAASFNKAMTNIVDAFTGVNDAITQTSQALQTVATALNKIQDVVNQQGNSLNHLT 848
Consensus         ** :***  :*   **.*:   :   .:  .               *:  *:*  .**  :..:*.  *

Heptad Repeat Region 1 End→←Central Helix Begin[1]                              Central
SARS2|QIK02964.   KQLSSNFGAISSVLNDILSRLDKVEAEVQIDRLITGRLQSLQTYVTQQLIRAAEIRASANLAA 1026
SARS|NP_828851.   KQLSSNFGAISSVLNDILSRLDKVEAEVQIDRLITGRLQSLQTYVTQQLIRAAEIRASANLAA 1008
MERS|QBM11748.1   SELSNTFGAISASIGDIIQRLDVLEQDAQIDRLINGRLTTLNAFVAQQLVRSESAALSAQLAK 1100
MHV-A59|AAU0635   NQLSNRFGAISASLQEILTRLEAVEAKAQIDRLINGRLTALNAYISKQLSDSTLIKVSAAQAI 1076
OC43|YP_0095552   QQLSNRFGAISASLQEILSRLDALEAEAQIDRLINGRLTALNAYVSQQLSDSTLVKFSAAQAM 1110
HKU1|Q14EB0.1     QQLFNKFGAISSSLQEILSRLDNLEAQVQIDRLINGRLTALNAYVSQQLSDITLIKAGASRAI 1107
NL63|YP_003767.   SQLRHNFQAISNSIQAIYDRLDSIQADQQVDRLITGRLAALNAFVSQVLNKYTEVRGSRRLAQ 1092
228E|NP_073551.   SQLRQNFQAISSSIQAIYDRLDTIQADQQVDRLITGRLAALNVFVSHTLTKYTEVRASRQLAQ 911
Consensus         .:*    * ***   :   *   **:  :: .  *:****.*** :*:.:::: *          .    *

 Helix End→                                                        ←Connector
SARS2|QIK02964.   TKMSECVLGQSKRVDFCGKGYHLMSFPQSAPHGVVFLHVTYVPAQEKNFTTAPAICHDGK--- 1086
SARS|NP_828851.   TKMSECVLGQSKRVDFCGKGYHLMSFPQAAPHGVVFLHVTYVPSQERNFTTAPAICHEGK--- 1068
MERS|QBM11748.1   DKVNECVKAQSKRSGFCGQGTHIVSFVVNAPNGLYFMHVGYYPSNHIEVVSAYGLCDSANPTN 1163
MHV-A59|AAU0635   EKVNECVKSQTTRINFCGNGNHILSLVQNAPYGLYFIHFSYVPISFTTANVSPGLCISGDR-- 1137
OC43|YP_0095552   EKVNECVKSQSSRINFCGNGNHIISLVQNAPYGLYFIHFSYVPTKYVTARVSPGLCIAGDR-- 1171
HKU1|Q14EB0.1     EKVNECVKSQSPRINFCGNGNHILSLVQNAPYGLLFIHFSYKPTSFKTVLVSPGLCLSGDR-- 1168
NL63|YP_003767.   QKINECVKSQSNRYGFCGNGTHIFSIVNSAPDGLLFLHTVLLPTDYKNVKAWSGICVDGIY-- 1153
228E|NP_073551.   QKVNECVKSQSKRYGFCGNGTHIFSIVNAAPEGLVFLHTVLLPTQYKDVEAWSGLCVDGTN-- 972
Consensus          *:.***  .*:  *  .***:*  *:.*:    ** *:  *:*     *  .          .:*   .

  Domain Begin[1]                                             Connector Domain End→
SARS2|QIK02964.   AHFPREGVFV--SNGT-----HWFVTQRNFYEPQIITTDNTFVSGNCDVVIGIVNNTVYDPLQ 1142
SARS|NP_828851.   AYFPREGVFV--FNGT-----SWFITQRNFFSPQIITTDNTFVSGNCDVVIGIINNTVYDPLQ 1124
MERS|QBM11748.1   CIAPVNGYFI--KTNNTRIVDEWSYTGSSFYAPEPITSLNTKYV-APQVTYQNISTNLPPPLL 1223
MHV-A59|AAU0635   GLAPKAGYFV--QDDG-----EWKFTGSSYYYPEPITDKNSVIMSSCAVNYTKAPEVFLNTSI 1193
OC43|YP_0095552   GIAPKSGYFV--NVNN-----TWMYTGSGYYYPEPITENNVVVMSTCAVNYTKAPYVMLNTSI 1227
```

#### Supplementary Material

```
HKU1|Q14EB0.1     GIAPKQGYFI--KQND-----SWMFTGSSYYYPEPISDKNVVFMNSCSVNFTKAPFIYLNNSI 1224
NL63|YP_003767.   GYVLRQPNLVLYSDNG-----VFRVTSRVMFQPRLPVLSDFVQIYNCNVTFVNISRVELHTVI 1211
228E|NP_073551.   GYVLRQPNLALYKEGN-----YYRITSRIMFEPRIPTMADFVQIENCNVTFVNISRSELQTIV 1030
Consensus                 :     .       :  *    : *.     :         *
```

←Heptad Repeat Region 2[3]

```
SARS2|QIK02964.   PE--LDSFKEELDKY-FKNHT-SPDVDLG-DISGINASVVNIQKEID--------------RL 1186
SARS|NP_828851.   PE--LDSFKEELDKY-FKNHT-SPDVDLG-DISGINASVVNIQKEID--------------RL 1168
MERS|QBM11748.1   GNSTGIDFQDELDEF-FKNVS-TSIPNFG-SLTQINTTLLDLTYEML--------------SL 1269
MHV-A59|AAU0635   PN--PPDFKEELDKW-FKNQT-SIAPDLSLDFEKLNVTLLDLTYEMN--------------RI 1238
OC43|YP_0095552   PN--LPDFKEELDQW-FKNQT-SVAPDLS--LDYINVTFLDLQVEMN--------------RL 1270
HKU1|Q14EB0.1     PN--LSDFEAEFSLW-FKNHT-SIAPNLTF-NSHINATFLDLYYEMN--------------VI 1268
NL63|YP_003767.   PD--YVDVNKTLQEF-AQNLPKYVKPNF--DLTPFNLTYLNLSSELKQLEAKTASLFQTTVEL 1269
228E|NP_073551.   PE--YIDVNKTLQELSYKL-PNYTVPDL--VVEQYNQTILNLTSEISTLENKSAELNYTVQKL 1088
Consensus          :    ..:  :.    :  .     ::        * : :::  *:                :
```

Heptad Repeat Region 2 End→←Transmembrane Domain[3]→← Cytoplasmic tail

```
SARS2|QIK02964.   NEVAKNLNESLIDLQELGKYEQYIKWPWYIWLGFIAGLIAIVMVTIMLCCMTSCCSC---LKG 1246
SARS|NP_828851.   NEVAKNLNESLIDLQELGKYEQYIKWPWYVWLGFIAGLIAIVMVTILLCCMTSCCSC---LKG 1228
MERS|QBM11748.1   QQVVKALNESYIDLKELGNYTYYNKWPWYIWLSFIAGLVALALCVFFILCCTGCGTN--CMG- 1329
MHV-A59|AAU0635   QDAIKKLNESYINLKEVGTYEMYVKWPWYVWLLIGLAGVAVCVLLFFICCCTGCGSC--CFK- 1298
OC43|YP_0095552   QEAIKVLNQSYINLKDIGTYEYYVKWPWYVWLLICLAGVAMLVLLFFICCCTGCGTS--CFK- 1330
HKU1|Q14EB0.1     QESIKSLNSSFINLKEIGTYEMYVKWPWYIWLLIVILFIIFLMILFFICCCTGCGSA--CFS- 1328
NL63|YP_003767.   QGLIDQINSTYVDLKLLNRFENYIKWPWWVWLIISVVFVVLLSLLVFCCLSTGCCGCCNCLT- 1331
228E|NP_073551.   QTLIDNINSTLVDLKWLNRVETYIKWPWWVWLCISVVLIFVVSMLLLCCCSTGCCGFFSCFA- 1150
Consensus         :   . :*.: ::*: :.    * ****::** :    : .    .:    *.*      :
```

ER or ERGIC localization signal[5]:← →

```
SARS2|QIK02964.   CCSCGSCCKFD--EDDSEP-VLKGVKLHYT 1273
SARS|NP_828851.   ACSCGSCCKFD--EDDSEP-VLKGVKLHYT 1255
MERS|QBM11748.1   KLKCNRCCDRY--EEYDLE--PH--KVHVH 1353
MHV-A59|AAU0635   --KCGNCCDEY--GGHQDSIVIHNISSHED 1324
OC43|YP_0095552   --KCGGCCDDY--TGYQE-LVIK--TSHDD 1353
HKU1|Q14EB0.1     --KCHNCCDEY--GGHND-FVIK--ASHDD 1351
NL63|YP_003767.   -SSMRGCCDCGSTKLPYYE--FE--KVHVQ 1356
228E|NP_073551.   -SSIRGCCEST--KLPYYD--VE--KIHIQ 1173
Consensus           .   **.             .
```

*Residues underlined have no coordinates in the structure file and are deemed flexible regions. The PDB files used to mark the same are SARS-CoV-2: 6VXX, SARS-CoV:5XLR, 6Q04: MERS-CoV, MHV-A59:3JCL, HCoV-OC43:6NZK, HCoV-HKU1:5I08. The sequences for HCoV-NL63 and HCoV-2289E are not marked because they belong to α-coronaviruses, whereas all others are from β-coronaviruses. The RED marked region spanning Heptad repeat region 2 C-terminal and trans-membrane domain N-terminal region is also called the aromatic domain or the fourth fusion peptide FP-IV.*

##### Accession number of Genome/Spike protein sequences used in this alignment

1: AY700211.1 / AAU06356.1 (Murine hepatitis virus strain A59)
2: KT029139.1 / QBM11748.1 (MERS-CoV/KOR/KNIH/002_05_2015)
3: NC_004718.3 / NP_828851.1 (SARS coronavirus)
4: MN985325.1 / QIK02964.1 (2019-nCoV/USA-WA1/2020)
5. NC_005831.2 /YP_003767.1 (Human Coronavirus NL63,)
6. NC_006213.1 / YP_009555241.1 (Human coronavirus OC43 strain ATCC VR-759)
7. NC_006577.2 / Q14EB0.1 (Human coronavirus HKU1)
8. NC_002645.1 / NP_073551.1 (Human coronavirus 229E)

##### References used to annotate the alignment:


1 Wrapp, D. *et al.* Cryo-EM structure of the 2019-nCoV spike in the prefusion conformation. *Science* **367**, 1260-1263 (2020).

2 Hofmann, H. & Pöhlmann, S. Cellular entry of the SARS coronavirus. *Trends Microbiol* **12**, 466-472 (2004).

3 Xia, S. *et al.* Inhibition of SARS-CoV-2 (previously 2019-nCoV) infection by a highly potent pan-coronavirus fusion inhibitor targeting its spike protein that harbors a high capacity to mediate membrane fusion. *Cell research*, 1-13 (2020).

4 Singh, M. *et al.* A proline insertion-deletion in the spike glycoprotein fusion peptide of mouse hepatitis virus strongly alters neuropathology. *J Biol Chem* 294, 8064-8087, DOI:10.1074/jbc.RA118.004418 (2019).
5 Sadasivan, J., Singh, M. & Sarma, J. D. Cytoplasmic tail of coronavirus spike protein has intracellular targeting signals. J Biosci 42, 231-244, DOI:10.1007/s12038-017-9676-7 (2017).

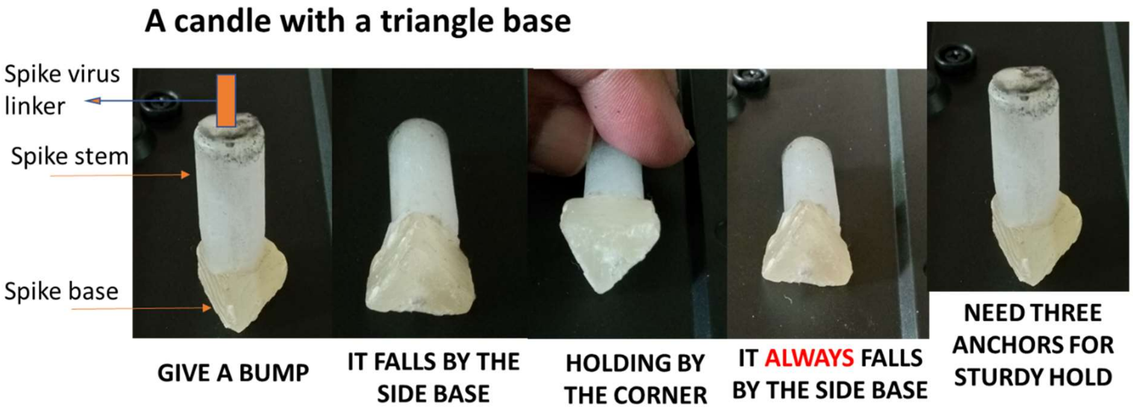


**Fig. S2.** A series of snapshots showing the properties of the Contact Initiation Model. The model is created using candle wax, where the base represents the triangular shape as found for the trimeric spike S1 domain, and the cylindrical stem represents the trimeric S2 domain. The stem is attached to the virus envelope through trimeric flexible linkers at the C-terminal end of the Spike ectodomain. The model on the extreme left shows that an unanchored structure on a triangle base, when bumped from the side, will fall in a position where one of the base sides rests on the surface. This in turn would ensure that the fusion peptide surface on the stem aligned along the middle of the base will make the initial contact and trigger the fusion process (Please check the triangles marked in Fig. 2D alongside). The fact that the orientation holding it by the corner of the base is not stable is emphasized in third and fourth illustrations, and the model always falls such that it rests on the side base. This architecture guarantees the fusion peptides in the S2 domain make the initial contact given that they are located at the crest of the bulge on the S2 domain surface. The last figure emphasizes that three anchors are needed for a stable vertical "tripod" orientation of the spike with respect to the host surface.

### Supplementary Material

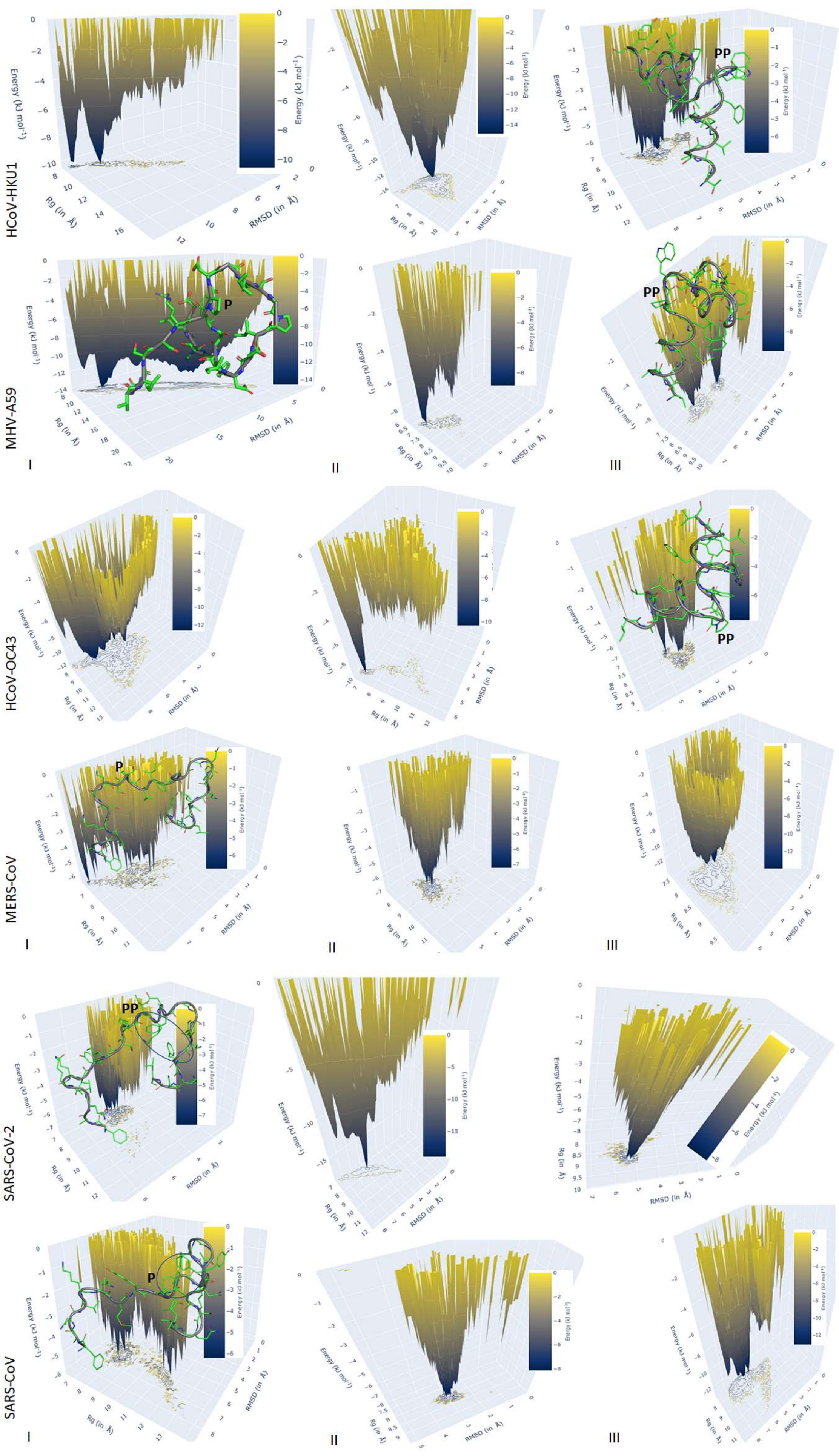

#### Supplementary Material

**Fig. S3. Energy landscape plot corresponding to all fusion peptides shown in Fig. 4 (main text)**. The ball and stick model of the starting structures of all the fusion peptides with central proline is shown along with their backbone trace. The central prolines are marked by “P”. Carbon atoms are in green, oxygen in red, nitrogen in blue, and sulfur in yellow. The location of the aromatic cluster is marked by oval in SARS-COV-2 and SARS-CoV Spike protein. The spread of the conformation in each FP can be visually estimated from the contours projected onto the X-Y plane.
